# Nuclear body assembly by a viral repeat RNA promotes Kaposi’s sarcoma-associated herpesvirus gene expression

**DOI:** 10.1101/2024.09.20.614208

**Authors:** Mariel Kleer, Rory P. Mulloy, Maxwell P. Bui-Marinos, Michael J. Johnston, Morgan F. Khan, David C. Schriemer, Jennifer A. Corcoran

**Affiliations:** Microbiology, Immunology and Infectious Diseases Department; Biochemistry and Molecular Biology Department; Arnie Charbonneau Cancer Institute; Robson DNA Science Centre; Snyder Institute for Chronic Diseases; Cumming School of Medicine, University of Calgary, Calgary, Alberta, Canada

## Abstract

*Kaposin* is the most abundantly expressed viral RNA in tumours caused by the oncogenic virus Kaposi’s sarcoma-associated herpesvirus (KSHV); however, its role in viral replication is not understood. Here, we show that *kaposin*, previously viewed as a protein-coding transcript, exists primarily as a nuclear viral long non-coding RNA (lncRNA) that rebuilds cellular nuclear speckles (NSs) adjacent to the viral genome to enhance viral gene expression. *Kaposin* is both necessary and sufficient to drive substantial NS remodelling, and this effect depends on repetitive elements within the RNA. Absence of *kaposin*-mediated NS remodeling, depletion of the essential NS protein, SRRM2, or blocking *kaposin* repetitive elements with antisense oligonucleotides impairs viral gene expression. *Kaposin* is the first viral lncRNA discovered to seed a functional nuclear body beside the viral genome. This work reframes our understanding of viral lncRNA function and the spatial organization of transcription in the infected cell nucleus.

## Introduction

Non-coding (nc) RNA serves as a molecular scaffold for the assembly of multiple nuclear compartments. This RNA-driven compartmentalization spatially organizes DNA, RNA, and proteins into “processing hubs” that regulate localized gene expression by concentrating effector proteins ^1–5^. One example of this is nuclear speckles (NSs) which are membraneless organelles (MLOs) composed of numerous proteins involved in RNA splicing and processing ^6–8^, a large fraction of polyadenylated RNA (poly(A)+ RNA) ^9^, and the long noncoding RNA (lncRNA) metastasis associated lung adenocarcinoma transcript 1 (*MALAT1*) ^9,10^. The assembly of NSs requires two large proteins, SON and serine/arginine repetitive matrix 2 (SRRM2), ^11^ and proceeds via initial homotypic SRRM2 interactions, followed by non-specific RNA binding, and culminating in phase separation ^12^. NSs regulate 3D genome organization, mRNA export, splicing, and transcription ^2,13–16^ by promoting RNA processing of genes in their spatial proximity. Genes localized near NSs are highly expressed, have low transcriptional pausing, and exhibit both increased spliceosome binding and increased co-transcriptional splicing ^15,16^. This makes NSs obvious targets for viruses that co-opt host processes to optimize viral gene expression. Indeed, many viruses modulate NSs ^17–25^; however, how this occurs and the resulting impact on viral and host gene expression remains largely unexplored.

Kaposi’s sarcoma-associated herpesvirus (KSHV) is a large double-stranded DNA virus that codes for at least 80 open reading frames (ORFs), several microRNAs, and small number of lncRNAs ^26^. KSHV is the causative agent of two lymphoproliferative disorders, primary effusion lymphoma and Multicentric Castleman’s disease, and the endothelial cell neoplasm Kaposi’s Sarcoma (KS) ^27–29^. The KSHV replication cycle can be divided into three phases: 1) primary infection 2) latency and 3) lytic reactivation. Primary infection includes initial viral entry and gene expression following which latency is established via ^35–37^ the physically tethering of the viral genome to the host chromosome by the viral latency associated nuclear antigen (LANA) ^30–32^ and transcriptional silencing save for a few select genes ^38–40^. KSHV remains in a latent state until cellular stress induces the expression of the viral replication and transcription activator (RTA) to initiate the lytic transcription cascade ^33–37^.

The viral transcript, *kaposin,* is expressed by all KS tumour cells regardless of tumour stage ^38,39^ and is one of the most abundant viral RNAs in all stages of KSHV infection ^26,40–42^. The exact length of the *kaposin* RNA varies between different KSHV strains ^40,41,43–45^. In the KSHV strain JSC-1, the most abundant version of the *kaposin* transcript is ∼1400 bp in size ^26,40^ and it contains two sets of GC-rich 23-nucleotide direct repeat (DR) regions termed DR2 (CACCCCAGGAACCCGGCGCGGCG) and DR1 (CCGGGAACCTGGTGCCCTCCTCC) that are repeated 21 and 27 times, respectively (Figure 1A). Shortly after its identification, *kaposin* was found to contain overlapping open reading frames encoding for three polypeptides: KapA, KapB and KapC (Figure 1A) ^46^. Since then, research from many labs, including our own, has focused on elucidating the function of these protein products ^47–52^.

**Figure 1.**
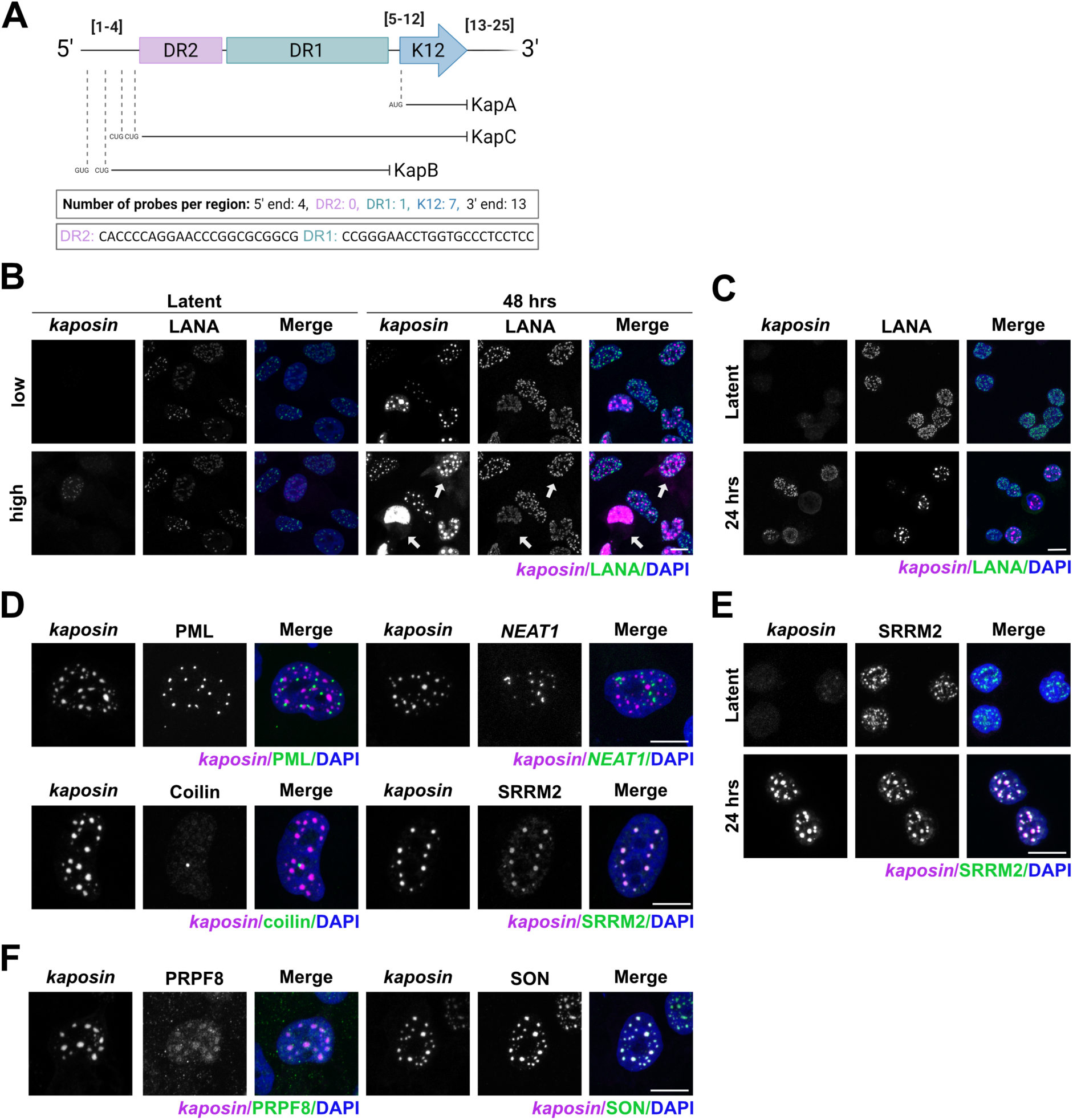
The *kaposin* RNA forms puncta that colocalize with nuclear speckles during KSHV infection. **(A)** The *kaposin* RNA, FISH probe annealing sites are indicated by numbers in square brackets. **(B)** Latent or reactivated iSLK- BAC16 (**B, D, F**) or BCBL-1 (**C, E**) cells were stained for *kaposin* and viral LANA (**B-C**) or *kaposin* and markers for nuclear bodies (**D, E, F**). Labels indicate: PML-PML bodies, Coilin-Cajal bodies, *NEAT1*- paraspeckles, SRRM2, PRPF8 or SON- nuclear speckles. White arrows indicate cytoplasmic *kaposin*.

To attribute function to individual kaposin polypeptides during infection, our lab used the KSHV bacterial artificial chromosome (BAC) system (BAC16; based on KSHV JSC-1) to build a panel of recombinant viruses deficient in kaposin protein expression individually and in combination^53^. Characterization of this panel revealed that all kaposin-recombinant viruses, regardless of the Kap protein(s) they lacked, were impaired relative to wild-type KSHV virus and these defects could not be rescued by the ectopic overexpression of *kaposin in trans*. This observation led us to hypothesize that optimal KSHV replication requires the *kaposin* RNA transcript *in cis* (at its site of transcription). We now reveal the mechanism that underlies this requirement; the *kaposin* transcript is a viral arcRNA that, upon transcription, recruits NS components to assemble NSs adjacent to the viral genome to optimize KSHV gene expression during all three phases of KSHV infection. Our work unveils a novel strategy used by herpesviruses to rearrange nuclear compartments within host nuclei to ensure successful infection and is the first example of an RNA transcript (cellular or viral) that is sufficient to seed NSs at a specific site, an event that serves to locally modulate gene expression.

## RESULTS

### *Kaposin* colocalizes with and remodels nuclear speckles

To investigate the role of the *kaposin* transcript we used the iSLK model of KSHV infection. iSLKs are a commonly used cell line in the KSHV field that harbor the doxycycline-inducible KSHV-encoded replication and transcription transactivator (RTA) transgene. After infection with Bacterial Artificial Chromosome (BAC)-derived KSHV (BAC16) ^54,55^, the addition of doxycycline and resulting RTA production is sufficient to transition the virus from latency into the lytic phase of infection, where full viral gene expression and progeny virion production will occur, a process called “reactivation”.

We hypothesized that the subcellular localization of *kaposin* (hereafter, italics denote the RNA transcript) may inform its function. To test this, we designed tiling FISH probes for the *kaposin* RNA (Figure 1A) and performed fluorescent *in situ* hybridization immunofluorescence (IF-FISH) for *kaposin* and the viral episome tethering protein LANA in iSLK-BAC16 cells during both latent and lytic (48 hrs post reactivation) infection. In latency only a small subset of cells exhibited visible *kaposin* staining, consistent with a low level of *kaposin* transcription during this replication phase in iSLKs ^26^. This contrasts with lytic reactivation, when most cells were intensely *kaposin*-positive, consistent with RTA-induced upregulation of *kaposin* transcription ^26,43^. Regardless of replication phase, *kaposin* localized mainly to the nucleus and exhibited a striking punctate staining pattern (Figure 1B). A small subset of cells undergoing lytic reactivation did exhibit cytoplasmic *kaposin* staining (Figure 1B, white arrows); however, these data suggest a large proportion of *kaposin* RNA is nuclear and unavailable for translation. We obtained similar results using a doxycycline-inducible B-cell model of latent and lytic KSHV infection derived from a primary effusion lymphoma (BCBLs) ^56^, underscoring that this staining pattern is not cell type specific (Figure 1C). These data suggest that *kaposin* may perform a previously unappreciated function as a non-coding RNA.

Intrigued by the punctate nature of *kaposin* staining, we asked whether these puncta colocalized with cellular nuclear bodies. To test this, we performed IF-FISH using lytic iSLK-BAC16s and co-stained for *kaposin* in combination with an established protein or RNA marker for four cellular nuclear bodies: PML bodies (promyelocytic leukemia protein, PML), Cajal bodies (coilin), paraspeckles (nuclear enriched abundant transcript 1, *NEAT1*), or nuclear speckles (SRRM2). While we observed minimal to no colocalization of *kaposin* with PML, coilin, or *NEAT1*, we found a striking colocalization of *kaposin* with the nuclear speckle (NS) marker SRRM2 (Figure 1D). Similarly, *kaposin* colocalized with SRRM2 in both latent and lytic BCBLs (Figure 1E). The strong colocalization we observed when we repeated the IF-FISH using two other markers of NSs, pre-mRNA processing factor 8 (PRPF8), and SON, (Figure 1F), indicated the *kaposin*-positive puncta we observed are not simply *kaposin*-SRRM2 aggregates.

NSs typically exhibit an irregular morphology ^57^; however, the *kaposin*-positive NSs we observed during reactivation were large and spherical, consistent with prior reports describing altered NS morphology during KSHV infection ^22,58^. To quantify these changes, we conducted a time course of KSHV infection in iSLK-BAC16s from latency to 72 hrs post reactivation and quantified the number, size, intensity, and eccentricity (non-roundness) of NSs (Figure 2A-B). We found that as lytic reactivation proceeded, NSs increased in size, intensity, and roundness, but decreased in overall number, presumably due to NS fusion ^59^. We also observed that host lncRNA, *MALAT1*, which normally localizes to the outer shell of NSs ^60^, was absent from NSs during KSHV reactivation (Figure 2C). Throughout reactivation *MALAT1-*SON colocalization decreased and *kaposin*-SON colocalization increased while *kaposin*-*MALAT1* colocalization was not observed (Figure 2D-E). To test whether *kaposin* expression alone is sufficient to induce NS remodeling, we overexpressed *kaposin* from a plasmid in the absence of infection. To ensure our observations were specific to the *kaposin* RNA, and not the Kap proteins, we created a construct in which all five previously identified *kaposin* translation initiation codons were eliminated (pcDNA-kaposinΔstarts), transfected increasing concentrations of this plasmid into cells, and performed IF-FISH for *kaposin* and SRRM2. We found that overexpression of *kaposinΔstarts* phenocopied the staining pattern of *kaposin* during infection by localizing predominantly to the nucleus and colocalizing with NS marker SRRM2 (Figure 2F). With increased *kaposin* abundance, we observed increased NS size, intensity, and number with no change to NS eccentricity (Figure 2G). Increasing *kaposin* expression via transfection was also sufficient to displace *MALAT1* from NSs (Figure 2H). Like during infection, *MALAT1-*SON colocalization decreased, *kaposin*-SON colocalization increased and *kaposin*-*MALAT1* colocalization was negligible (Figure 2I-J). We did not explore whether *MALAT1* and *kaposin* compete for binding of NS proteins as their sequence and sub-NS localization differ ^60,61^, making this unlikely. Together these data reveal that *kaposin* expression alone, independent of any other viral genes, is sufficient to colocalize with and remodel NS size, intensity, and RNA composition.

**Figure 2.**
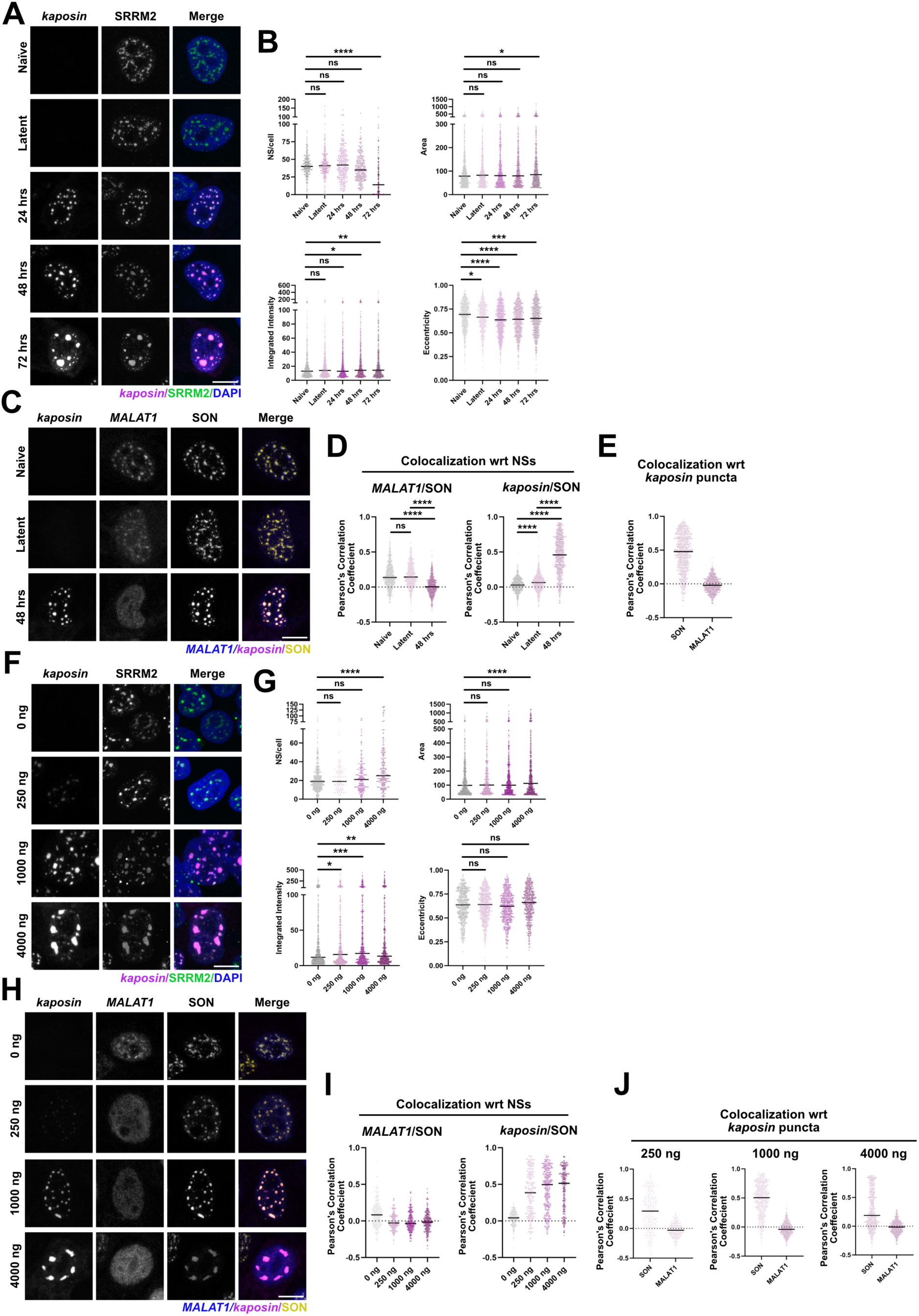
*Kaposin* RNA expression induces NS remodeling. **(A-E)** Naïve iSLK or iSLK-BAC16 cells were either not treated (naïve, latent) or reactivated for 24, 48, and 72 h and stained for *kaposin*, SRRM2 and nuclei (**A-B**) or *kaposin*, *MALAT1*, and SON (**C-E**). (**F-J**) Transfected 293T (**F-G**) or HeLa (**H-J**) cells were stained for *kaposin*, SRRM2, and nuclei (**F-G**) or *kaposin*, *MALAT1*, and SON (**H-J**). In B, D-E, G, and I-J, images were quantified using CellProfiler. A singular dot is equivalent to one NS (**B, D, G, I**) or one *kaposin*-positive foci (**E, J**). One-way ANOVA with a Tukey’s multiple comparisons post-hoc tests were performed (*, P < 0.05; ** P <0.01; *** P < 0.001; **** P < 0.0001; ns, nonsignificant) for all analyses. All experiments N=3 except **H-J**, N=2.

### *Kaposin* repeat sequences and cellular SRRM2 are required for *kaposin*-NS colocalization

The *kaposin* RNA contains two highly GC-rich direct repeat regions termed DR2 and DR1. While there is no known targeting sequence for NS localization, RNAs with repetitive sequences have previously been shown to be detained in NSs ^62^. To test whether the *kaposin* DRs mediate NS-colocalization we utilized our panel of recombinant viruses in which kaposin is either lacking DR1 and DR2 completely (BAC16ΔKapABC, BAC16ΔKapBC) or has been significantly recoded such that the DR regions are less repetitive and less GC-rich (BAC16ΔKapB, BAC16ΔKapC) ^53^. These viruses display replication defects compared to wild-type (WT) BAC16^53^. We also created an additional recombinant virus containing three 23-nucleotide repeats of DR2 and five 23-nucleotide repeats of DR1, called BAC16-miniDRs (Figure 3A, Table 1). We reactivated iSLKs harbouring each of the recombinant viruses in their latent form to enter lytic replication and performed IF-FISH for *kaposin* and SRRM2. During WT KSHV reactivation, *kaposin* colocalized with NS marker SRRM2, but in iSLK-BAC16 cells infected with kaposin-recombinant viruses (ΔKapABC, ΔKapBC, ΔKapB and ΔKapC), colocalization with SRRM2 was not observed (Figure 3B). iSLK-BAC16-miniDRs infected cells displayed a mixed phenotype where partial *kaposin*-SRRM2 colocalization was observed (Figure 3B). WT, ΔKapB and ΔKapC-reactivated iSLK cells demonstrated Kap protein translation although *kaposin* transcript copy number was decreased during ΔKapB, ΔKapC, and miniDR reactivation (Supp Figure 1A-B). Biochemical fractionation and RT-qPCR confirmed the majority of WT *kaposin* was nuclear-localized whereas the recoded *kaposin* transcripts expressed by ΔKapB and ΔKapC viruses (lacking both DRs) exhibited increased cytoplasmic localization (Supp Figure 1C-D), consistent with our earlier IF-FISH findings. Taken together, these results reveal that wild-type *kaposin* DR sequences are required for *kaposin*-SRRM2 colocalization and nuclear retention during infection and that three copies of DR2 and five copies of DR1 provide sufficient signal to partially rescue this colocalization.

**Figure 3.**
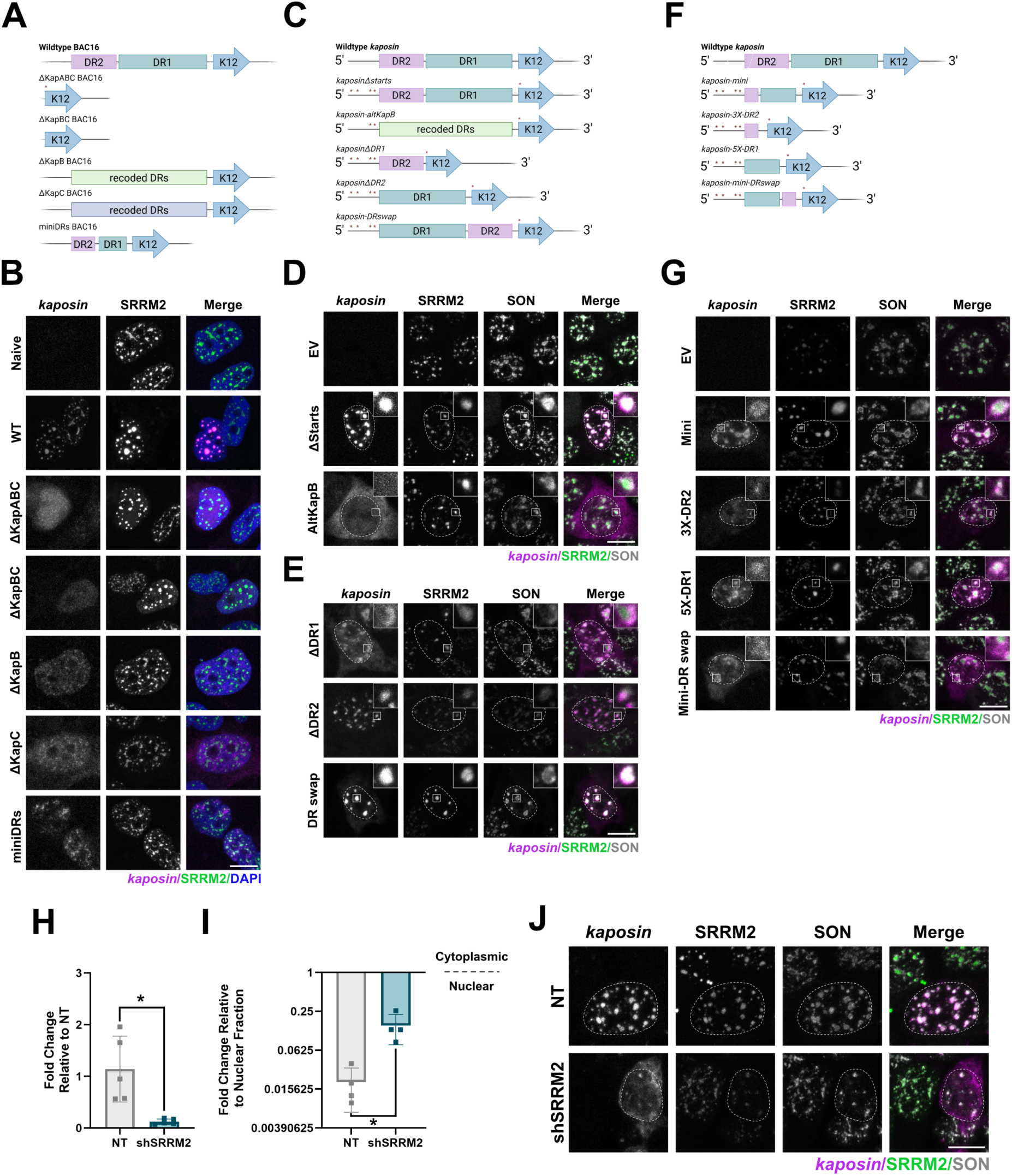
*Kaposin* repeat sequences and cellular SRRM2 are required for *kaposin*-NS colocalization. **(A, C, F)** The *kaposin* RNA as expressed from recombinant BAC16 viruses (**A,** Table 1) or pcDNA3 plasmids (**C, F;** Tables 2, 3 ) are shown. A red asterisk = mutated start codon. **(B)** Naïve or infected BAC16 iSLK cells were reactivated for 48h and stained for *kaposin*, SRRM2, and nuclei. Colour threshold for *kaposin* channel in WT iSLK condition for visualization purposes only. **(D-E, G)** 293T cells were transfected as indicated and stained for *kaposin*, SRRM2, and SON. **(H-J)** 293T cells transduced with an shRNA targeting SRRM2 lentiviruses were lysed to confirm knockdown by RT-qPCR with *SRRM2* or *18S* (housekeeping) specific primers (paired Student’s t-test (*, P < 0.05; ** P <0.01; *** P < 0.001; **** P < 0.0001; ns, nonsignificant). N=5**) (G)** or transfected with pcDNA-kaposinΔstarts **(H-I)**. Transfected cells were either fractionated, RNA extracted and RT-qPCR performed using *kaposin* and *18S* (housekeeping) specific primers (paired Student’s t- test (*, P < 0.05; ** P <0.01; *** P < 0.001; **** P < 0.0001; ns, nonsignificant)). N=4 **(H)** or fixed for IF-FISH and stained for *kaposin*, SRRM2 and SON (**I**). Dashed white outline indicates nucleus.

**Table 1.**
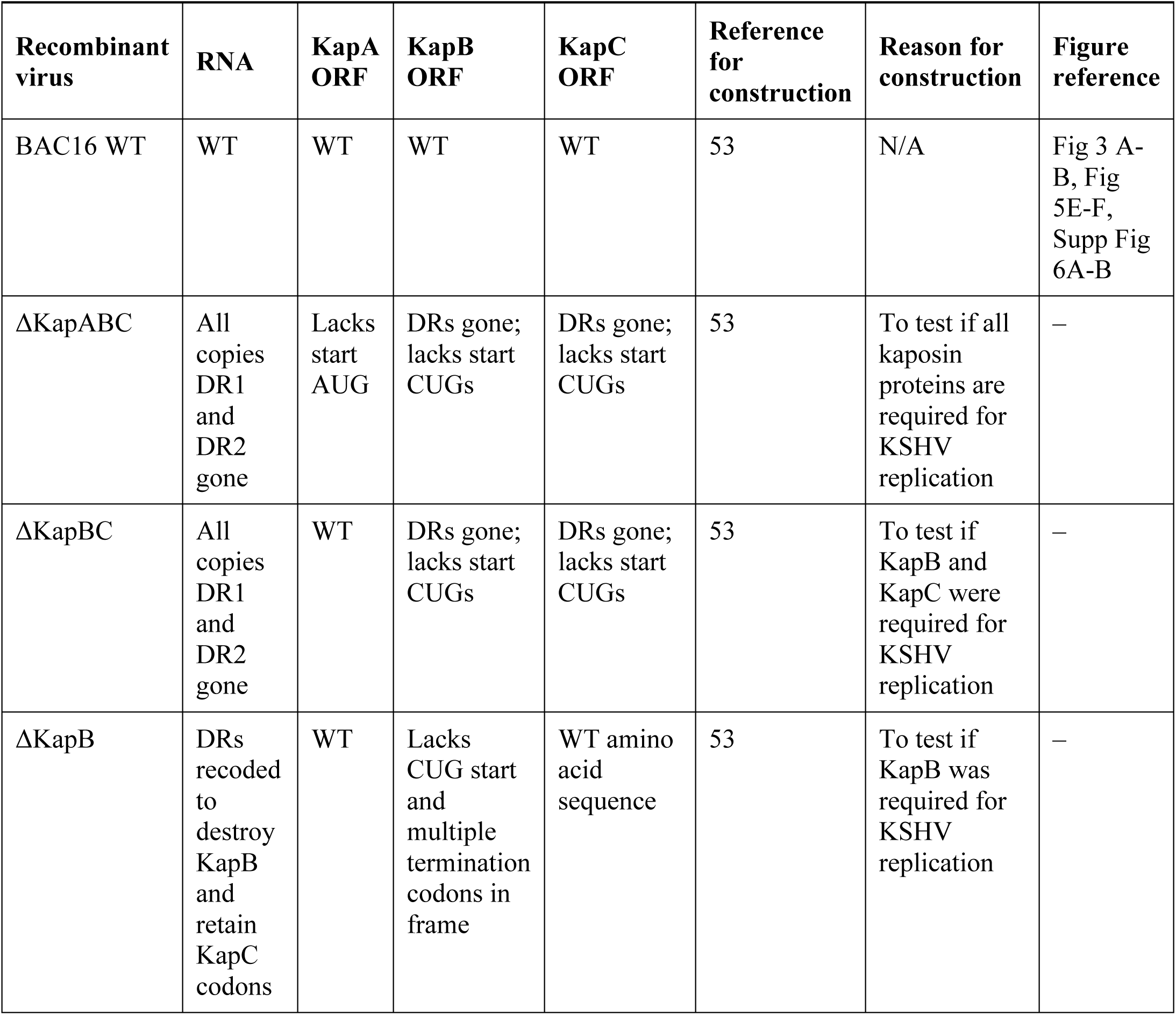
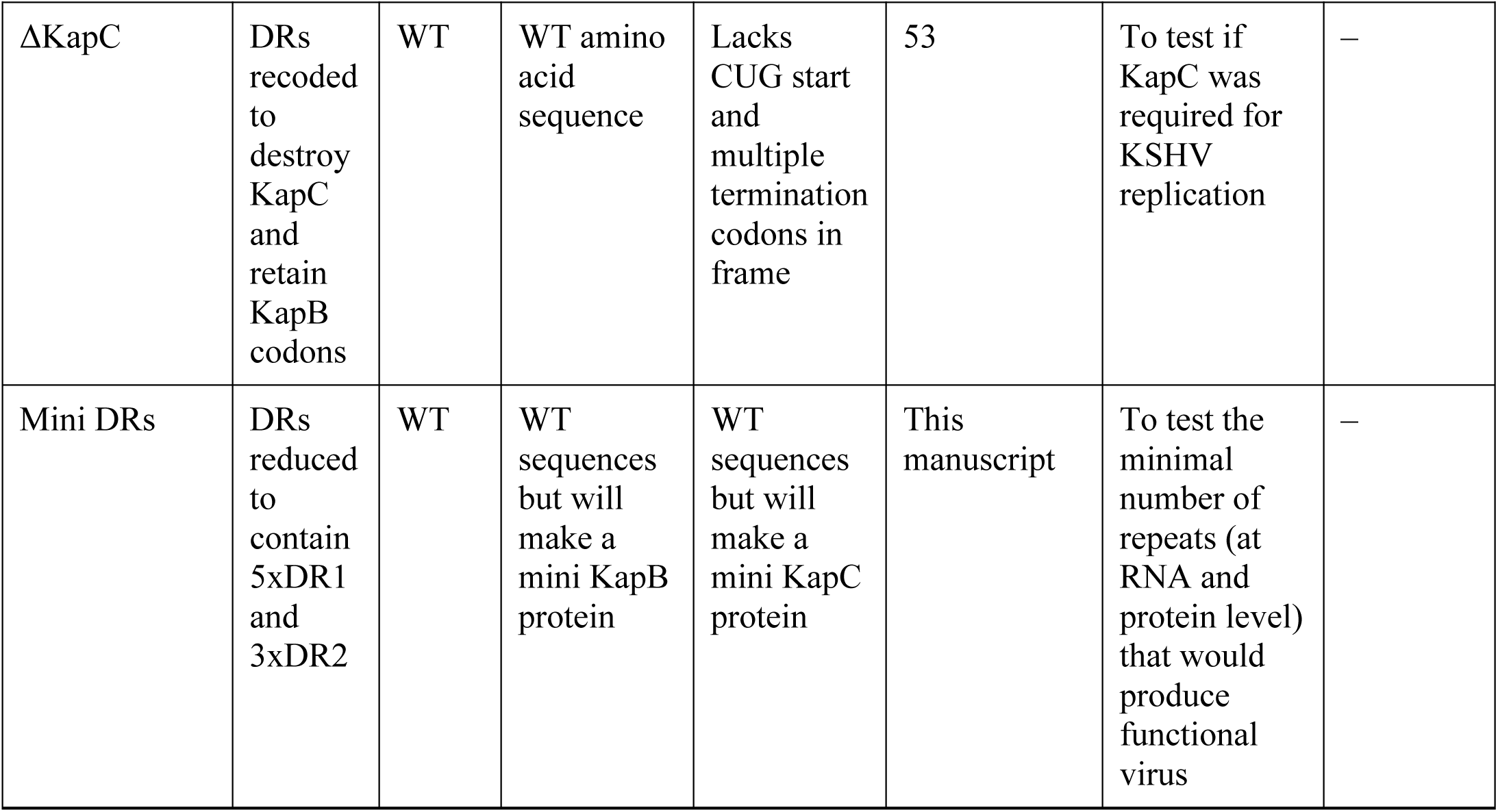
Recombinant Viruses Related to Figure 3A–B, Figure 5E-F and Supp Figure 6A–B.

**Table 2.**
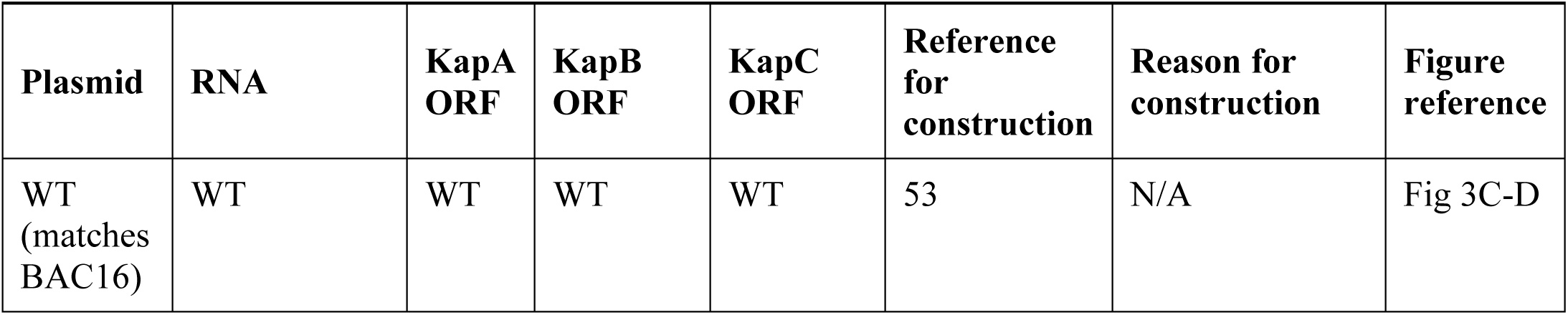
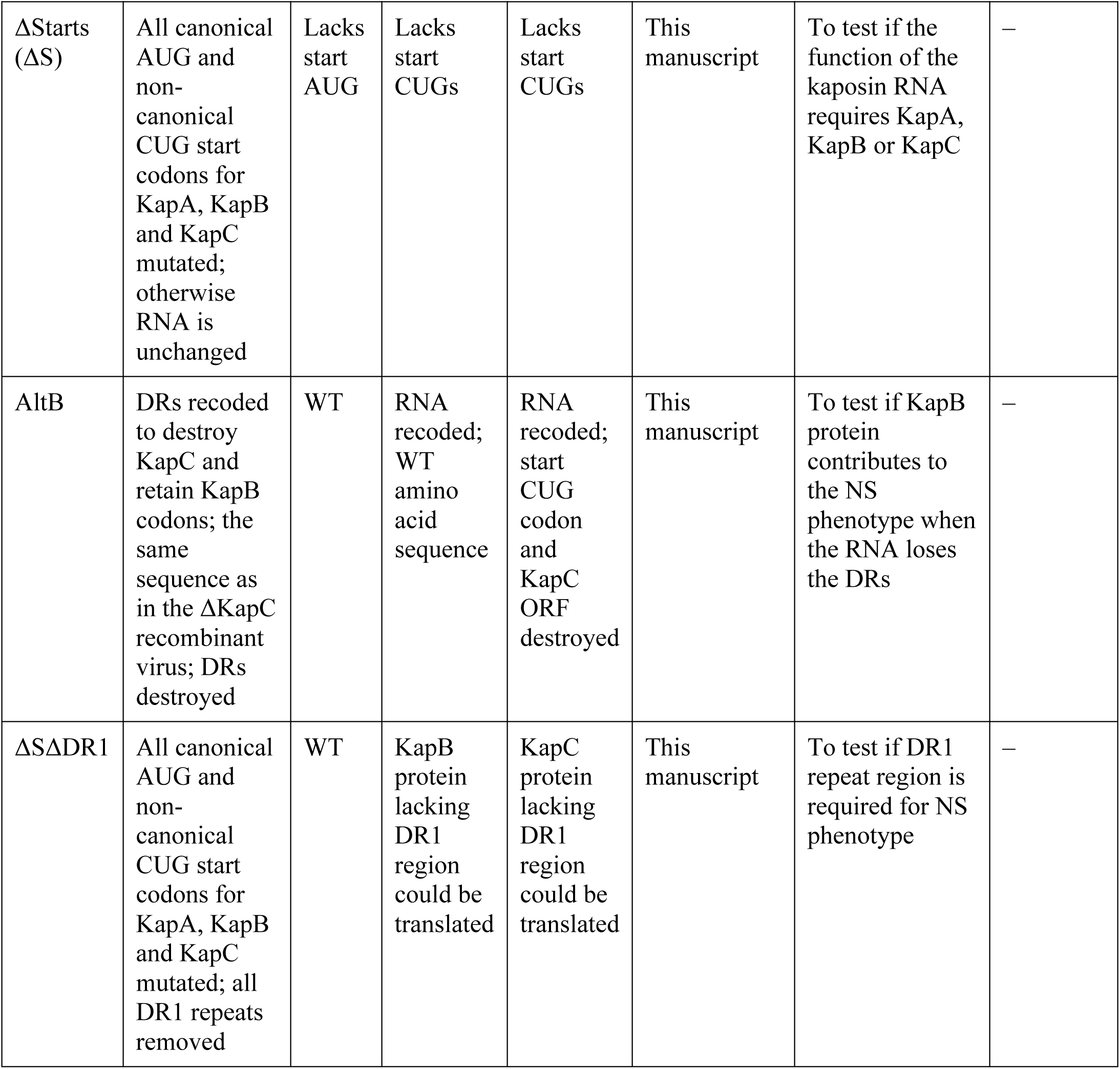
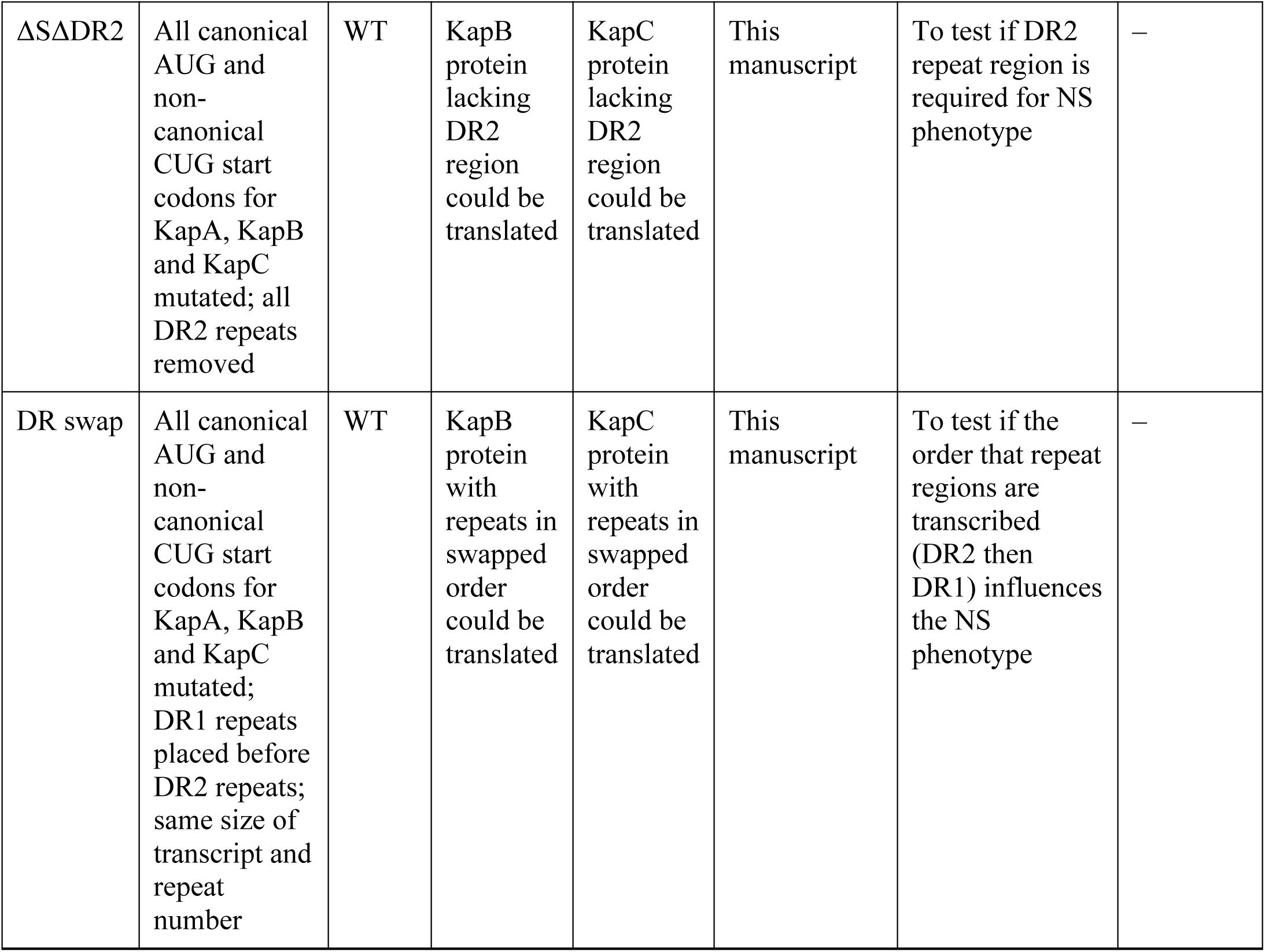
Plasmid constructs Related to **Figure 3C–D**.

| **Plasmid** | **RNA** | **KapA ORF** | **KapB ORF** | **KapC ORF** | **Reference for construction** | **Reason for construction** | **Figure reference** |
| --- | --- | --- | --- | --- | --- | --- | --- |
| WT (matches BAC16) | WT | WT | WT | WT | 53 | N/A | Fig 3C-D |

| **Plasmid** | **RNA** | **KapA**<br>**ORF** | **KapB**<br>**ORF** | **KapC**<br>**ORF** | **Reference**<br>**for**<br>**construction** | **Reason for**<br>**construction** | **Figure**<br>**reference** |
| --- | --- | --- | --- | --- | --- | --- | --- |
| ΔStarts<br>(ΔS) | All canonical AUG and non-canonical CUG start codons for KapA, KapB and KapC mutated; otherwise RNA is unchanged | Lacks start AUG | Lacks start CUGs | Lacks start CUGs | This manuscript | To test if the function of the kaposin RNA requires KapA, KapB or KapC | — |
| AltB | DRs recoded to destroy KapC and retain KapB codons; the same sequence as in the ΔKapC recombinant virus; DRs destroyed | WT | RNA recoded; WT amino acid sequence | RNA recoded; start CUG codon and KapC ORF destroyed | This manuscript | To test if KapB protein contributes to the NS phenotype when the RNA loses the DRs | — |
| ΔSΔDR1 | All canonical AUG and non-canonical CUG start codons for KapA, KapB and KapC mutated; all DR1 repeats removed | WT | KapB protein lacking DR1 region could be translated | KapC protein lacking DR1 region could be translated | This manuscript | To test if DR1 repeat region is required for NS phenotype | — |
| $\Delta S\Delta DR2$ | All canonical AUG and non-canonical CUG start codons for KapA, KapB and KapC mutated; all DR2 repeats removed | WT | KapB protein lacking DR2 region could be translated | KapC protein lacking DR2 region could be translated | This manuscript | To test if DR2 repeat region is required for NS phenotype | – |
| DR swap | All canonical AUG and non-canonical CUG start codons for KapA, KapB and KapC mutated; DR1 repeats placed before DR2 repeats; same size of transcript and repeat number | WT | KapB protein with repeats in swapped order could be translated | KapC protein with repeats in swapped order could be translated | This manuscript | To test if the order that repeat regions are transcribed (DR2 then DR1) influences the NS phenotype | – |

**Table 3.** Plasmid Constructs Related to **Figure 3E–F**.

| **Plasmid** | **RNA** | **KapA**<br>**ORF** | **KapB**<br>**ORF** | **KapC**<br>**ORF** | **Reference**<br>**for**<br>**construction** | **Reason for**<br>**construction** | **Figure**<br>**reference** |
| --- | --- | --- | --- | --- | --- | --- | --- |
| WT<br>(matches<br>BAC16) | WT | WT | WT | WT | 53 | N/A | Fig 3E-F |
| Mini | WT<br>sequence;<br>minimal<br>number of<br>DR1 (5x)<br>and DR2<br>(3x) repeats;<br>designed to<br>maintain the<br>ratio of<br>DR1:DR2<br>repeat<br>regions<br>(always more<br>DR1 than<br>DR2) | WT | WT<br>KapB<br>protein<br>with<br>shorter<br>stretches<br>of both<br>amino<br>acid<br>repeats<br>could be<br>translated | WT<br>KapC<br>protein<br>with<br>shorter<br>stretches<br>of both<br>amino<br>acid<br>repeats<br>could be<br>translated | This<br>manuscript | To determine the<br>minimal repeat<br>number of DR1<br>plus DR2 that<br>retains the NS<br>phenotype | — |
| 3xDR2 | WT<br>sequence<br>surrounding<br>the DR2<br>repeat<br>region;<br>minimal<br>number of<br>DR2 repeats | WT | KapB<br>protein<br>with<br>shorter<br>stretch of<br>DR2<br>amino<br>acid<br>repeats<br>lacking<br>DR1<br>repeats<br>could be<br>translated | KapC<br>protein<br>with<br>shorter<br>stretch of<br>DR2<br>amino<br>acid<br>repeats<br>lacking<br>DR1<br>repeats<br>could be<br>translated | This<br>manuscript | To determine the<br>minimal number<br>of DR2 repeats<br>that are required<br>to exhibit partial<br>NS<br>colocalization<br>(SON only) | — |
| 5xDR1 | WT<br>sequence<br>surrounding<br>the DR1<br>repeat<br>region;<br>minimal<br>number of<br>DR1 repeats | WT | KapB<br>protein<br>with<br>shorter<br>stretch of<br>DR1<br>amino<br>acid<br>repeats<br>lacking<br>DR2<br>repeats<br>could be<br>translated | KapC<br>protein<br>with<br>shorter<br>stretch of<br>DR1<br>amino<br>acid<br>repeats<br>lacking<br>DR2<br>repeats<br>could be<br>translated | This<br>manuscript | To determine the<br>minimal number<br>of DR2 repeats<br>that are required<br>to exhibit<br>NS/SRRM2<br>colocalization | — |
| Mini-<br>swap | All canonical<br>AUG and<br>non-canonical<br>CUG start<br>codons for<br>KapA, KapB<br>and KapC<br>mutated; all<br>DR2 repeats<br>removed | WT | KapB<br>protein<br>with<br>minimal<br>DR<br>repeats in<br>swapped<br>order<br>could be<br>translated | KapC<br>protein<br>with<br>minimal<br>DR<br>repeats in<br>swapped<br>order<br>could be<br>translated | This<br>manuscript | To test if the<br>order that repeat<br>regions are<br>transcribed<br>(DR2 then DR1)<br>influences the<br>NS phenotype in<br>the context of<br>the minimal DR<br>repeat construct | — |

To reveal which aspect of the *kaposin* DR sequences is required for NS colocalization, we designed several *kaposin* plasmid constructs for transfection. In addition to *kaposinΔstarts* construct, this included i) alternatively coded KapB (*kaposin-altKapB*), where the DR sequence is significantly reduced in its GC-rich and repetitive nature, but the mRNA still encodes the KapB protein (same RNA as ΔKapC) ii) *kaposinΔDR1*, where the DR1 repeat sequence is completely removed but the flanking nucleotide sequences and DR2 repeats are unaltered iii) *kaposinΔDR2*, where the DR2 repeat sequence is completely removed but the flanking nucleotide sequences and DR1 repeats are unaltered and iv) *kaposin-DRswap*, where the order of the DRs is switched such that DR1 is located 5’ to DR2 instead of 3’ to DR2 (Figure 3C, Table 2). We transfected each of these constructs into cells and performed IF-FISH for *kaposin* and SRRM2 and SON which localize to the inner and outer shell of the NS, respectively ^60^ (Figure 3D). Two constructs (*kaposinΔstarts* and *kaposinΔDR2*) colocalized with both NS markers whereas recoding the DRs (*kaposin-altKapB)* abolished NS colocalization and resulted in an increase in cytoplasmic localization of the *kaposin-altKapB* RNA (Figure 3D, Suppl Figure 2D). Cells expressing *kaposinΔDR1* exhibited a strong increase in cytoplasmic *kaposin* staining coupled with a loss of SRRM2 colocalization; however, a fraction of the *kaposin* in these cells remained in the nucleus and colocalized with SON in a donut-shaped pattern peripheral to the SRRM2 core of the NS. Taken together, these data support a model where the DR1 region is required for NS colocalization via a direct or indirect interaction with the core NS protein, SRRM2. Because *kaposinΔDR1* colocalized with SON, whereas *kaposin-altKapB* did not, we speculate that DR2 may provide a secondary NS localization signal via direct or indirect interaction with SON. Our observations suggest that of these two localization signals, DR1-mediated SRRM2 recruitment dominates, rendering the DR2-mediated SON recruitment redundant unless DR1 is removed. To assess whether the 5’-3’ sequence orientation of *kaposin* is likewise required for stepwise *kaposin*-mediated NS assembly, we overexpressed a *kaposin-DRswap* construct, in which DR1 is transcribed prior to DR2. Swapping the order of the DRs resulted in a small increase in cytoplasmic *kaposin* staining, yet a large fraction of *kaposin* staining still colocalized with SRRM2 and SON (Figure 3D-E), suggesting that the order of arrangement of the repeat regions of full-length *kaposin* is not essential.

The repetitive nature of *kaposin* (21 copies of DR2 followed by 27 copies of DR1) suggests that each repeat unit may function as an iterative signal for individual NS protein recruitment. To determine if the number of DRs effects NS colocalization we created a series of constructs based on the *kaposin-miniDR* background (Figure 3F, Table 3). *Kaposin-miniDR* encodes three DR2 repeats followed by five DR1 repeats and matches the *kaposin* nucleotide sequence of our miniDRs virus. We expressed this construct and two constructs containing each mini DR on its own (*kaposin-3X-DR2* and *kaposin-5X-DR1,* Table 3) and found that *kaposin-miniDR* colocalized with both SRRM2 and SON; however, this colocalization was weak relative to *kaposinΔstarts* and was coupled with a small amount of cytoplasmic *kaposin* staining (Figure 3F). Cells expressing *kaposin-3X-DR2* exhibited a small amount of colocalization with SON, whereas cells expressing *kaposin-5X-DR1* exhibited weak *kaposin* colocalization with both NS markers. Finally, curious whether a lowered NS-colocalization affinity would reveal a greater dependence on DR order, we created a construct where 5X-DR1 is transcribed prior to 3X-DR2 (*kaposin-mini-DRswap*) (Figure 3F-G, Table 3). Swapping the order of the mini DRs resulted in a stark increase in cytoplasmic *kaposin* staining with only minor NS colocalization, mostly with SON. These data suggest three things. i) Increased DRs repeats correspond to increased NS-colocalization, indicating that the DRs are iterative NS-colocalization signals. ii) Both DRs provide NS localization signals as the colocalization of either alone is weaker than that of the combined construct. iii) The placement of DR1 3’ to DR2 favours *kaposin-*NS colocalization and this requirement can be overcome through additional repeats. Based on this series of observations, we speculate that SON and SRRM2 interact with *kaposin* in a sequential, co-transcriptional, and iterative manner and that in the absence of SON and SRRM2 recruitment, *kaposin* is not retained in NSs and is instead available to be exported. After transfection, several constructs retained some ability to translate Kap proteins, including *kaposinΔstarts* which lacks the start codons reported for KapA, KapB and KapC, suggesting alternative translation mechanisms^63,64^. Probe-based TaqMan PCR revealed the transcript copy number was not significantly different between constructs indicating that expression level was not the reason for differential localization of transcripts (Supp Figure 2A-B). Biochemical fractionation followed by transcript copy number analysis recapitulated *kaposin* localization via FISH, with most *kaposin* transcripts exhibiting strong nuclear localization except for *kaposin-altKapB*. The cytoplasmic fraction of *kaposinΔDR1* observed by FISH was reduced in these experiments (Supp Figure 2C-D).

To test whether SRRM2 abundance alters *kaposin*-NS colocalization, we transfected either NT-control or SRRM2-silenced cells with *kaposinΔstarts,* performed nuclear-cytoplasmic fractionation for total RNA, and quantified the abundance of *kaposin* RNA in each fraction. These data are presented as a cytoplasmic:nuclear ration where values >1 indicate cytoplasmic localization whereas values <1 indicate nuclear localization. Relative to control, in SRRM2-silenced cells, there was a significant increase in cytoplasmic *kaposin* RNA localization (Figure 3H-I). In parallel, we observed increased *kaposin* RNA signal in the cytoplasm of SRRM2-silenced cells by IF-FISH (Figure 3J). These data reveal that *kaposin*-NS colocalization is mediated at least in part by SRRM2 and that decreased SRRM2 expression diminishes *kaposin* nuclear retention.^77,78^

### *kaposin*-NS colocalization does not require a linear sequence motif

RNAs with 5’ splice sites (SS), SR binding motifs, and hairpin secondary structures have all been shown to localize to NS ^62,65^. To determine whether a specific linear sequence motif or RNA secondary structure is required for *kaposin*-NS colocalization, we generated *kaposin-reverse*, a construct with the *kaposin* nucleotide sequence in reverse (3’->5’) orientation (Supp Figure 3A, Table 5). Following expression, we found that *kaposin-reverse* colocalized with both NS markers, SON and SRRM2, although the intensity of staining was greatly diminished suggesting decreased expression (Supp Figure 3B). These data suggest that *kaposin* localization to NSs can occur independent of a 5’->3’ sequence motif.

To confirm our findings using an orthogonal approach we used the MS2-MCP system ^66^ to visualize *kaposin* subcellular localization in a probe-independent manner. We designed a series of MS2-tagged constructs that included i) an “EV” control containing the MS2 sequences only, *EV-MS2* ii) a full length kaposinΔstarts construct, *kaposinΔstarts-MS2*, iii) a construct containing just four repeats of DR2, *kaposin-4X-DR2-MS2* and iv) a construct containing just four repeats of DR1, *kaposin-4X-DR1-MS2* (Supp Figure 3C, Table 5). Upon transfection into 293T cells, we found that overexpression of *EV-MS2* led to a largely diffuse pattern of MCP localization with many cells exhibiting very small puncta that did not colocalize with either SON or SRRM2 (Supp Figure 3D). In cells overexpressing *kaposinΔstarts-MS2,* there was a drastic reorganization of MCP staining into large nuclear bodies that colocalized with both SON and SRRM2, recapitulating the pattern of *kaposin* localization observed in Figures 2 and 3. After overexpression of *kaposin-4X-DR2-MS2,* MCP-positive foci co-localized with SON but not SRRM2 in a subset of cells, mirroring the staining pattern of *kaposinΔDR1*. By contrast, after *kaposin-4X-DR1-MS2* overexpression, MCP did not colocalize with either NS marker, like that of the *EV-MS2* control (Supp Figure 3D). These results indicate that i) *kaposin*-NS colocalization can be detected in a probe-independent manner, ii) MS2 stem loops alone are not sufficient to mediate NS colocalization, and iii) four repeats of DR2 are sufficient to mediate SON colocalization in a subset of cells, but four repeats of DR1 are insufficient to mediate SON or SRRM2 colocalization in this assay. Next, we asked whether this was occurring in a sequence orientation or GC-content dependent manner. To test this, we used *EV-MS2*, *kaposin-4X-DR2-MS2* and two additional constructs: one with the DR2 sequence in the reverse orientation (3’- >5’), called *kaposin-4X-DR2-reverse-MS2*, and one with the DR2 repeat scrambled, preserving GC-content but altering the repeat sequence, called *kaposin-4X-DR2-scrambled-MS2* (Supp Figure 3E, Table 5). Transfection and expression of these constructs revealed that while *kaposin-4X-DR2-reverse-MS2* still colocalized with SON, colocalization of *kaposin-4X-DR2-scrambled-MS2* with SON was markedly reduced (Supp Figure 3F).

Collectively, these data show that neither a linear sequence motif nor GC-content drive *kaposin* co-localization with NSs, suggesting this effect may instead be mediated by RNA structure. To this end, we used RNAstructure ^67^ to predict the secondary structure of the *4X-DR2* and *4X-DR1* RNAs. In both cases, large regions of double-stranded RNA were predicted (Supp Figure 3G). For *4X-DR1*, two stem loop structures were predicted for which the base pairing was highly probable. Rhesus macaque rhadinovirus (RRV) is a related herpesvirus that encodes an RNA transcript that is also GC-rich and repetitive. It contains two sets of direct repeats termed rDL E2 (32 bp repeat, repeated 32 times) and rDL E1 (19 bp repeat, repeated 21 times) ^68^ an organization that mirrors that of *kaposin*. To examine whether the repeats from RRV also confer NS colocalization, we inserted either three copies of the RRV rDL E2 repeat or five copies of the RRV rDL E1 repeat into our backbone EV-MS2 construct. We transfected these constructs alongside controls and found that in overexpression of RRV *3X-rDL-E2-MS2,* MCP co-localized with SON while in overexpression of *RRV 5X-rDL-E1-MS2,* the MCP signal remained diffuse (Supplemental Figure 3H). These data indicate that three copies of RRV rDL E2 repeats are sufficient to confer SON-colocalization, analogous to overexpression of *kaposin-4X-DR2* and suggests that repetitive viral RNA-mediated NS colocalization may be a herpesvirus conserved strategy.

### *Kaposin* transcription seeds a NS *de novo*

*Kaposin* may i) be a passenger RNA that is transcribed and subsequently transported to a pre-formed NS or ii) a driver RNA scaffold that recruits NS proteins to its site of transcription, inducing the formation of a NS *de novo*. To address this, we conducted live-cell microscopy using the MS2-MCP system by co-transfection of the following components: a doxycycline-inducible *kaposinΔstarts-MS2* construct, fluorescently tagged MCP, and two fluorescently tagged NS-resident proteins, SRSF1 and RMB25 (Figure 4A). We found that MCP localization was not altered by transfection and Dox-induction of *EV-MS2* alone (Video 1); however, in cells transfected with the *kaposinΔstarts-MS2* construct, MCP re-localized into large puncta over the two-hour period following dox-induced transcription (Video 2). Small MCP-positive foci were observed a few minutes post Dox addition, likely indicating the initial sites of *kaposin* transcription. *Kaposin* nascent RNA foci remained at their initial site of appearance, did not relocate, and became enriched for NS proteins either almost immediately (RMB25, Figure 4B, Video 3) or several minutes later (SRSF1, Figure 4C, Video 4). Over the duration of imaging, *kaposin* nascent RNA foci grew and became more spherical in nature (Video 5). These data show that nascent *kaposin* recruits NS proteins to its site of transcription and suggest *kaposin* drives NS assembly *de novo.* Once formed, *kaposin*-positive NS are dynamic in nature, and display features consistent with a phase-separated granule, including fusion and fission events (Video 5). To corroborate these findings using core NS markers SON and SRRM2, we transfected cells with Dox-inducible *EV-MS2*, *kaposinΔstarts-MS2*, *kaposinΔDR1-MS2*, and *kaposinΔDR2-MS2* expression constructs (Table 4), and fixed at 0, 30 min, 1 hr, and 2 hr timepoints post-dox induction. In cells expressing the *EV-MS2* control, MCP puncta were small and seldom colocalized with either SON or SRRM2 (Figure 4D). By contrast, in cells expressing *kaposinΔstarts-MS2*, MCP colocalized with both SON and SRRM2 at all time points. These foci were initially small at 30 min post-dox but were larger at both 1 and 2 hr post-dox (Figure 4D).

**Figure 4.**
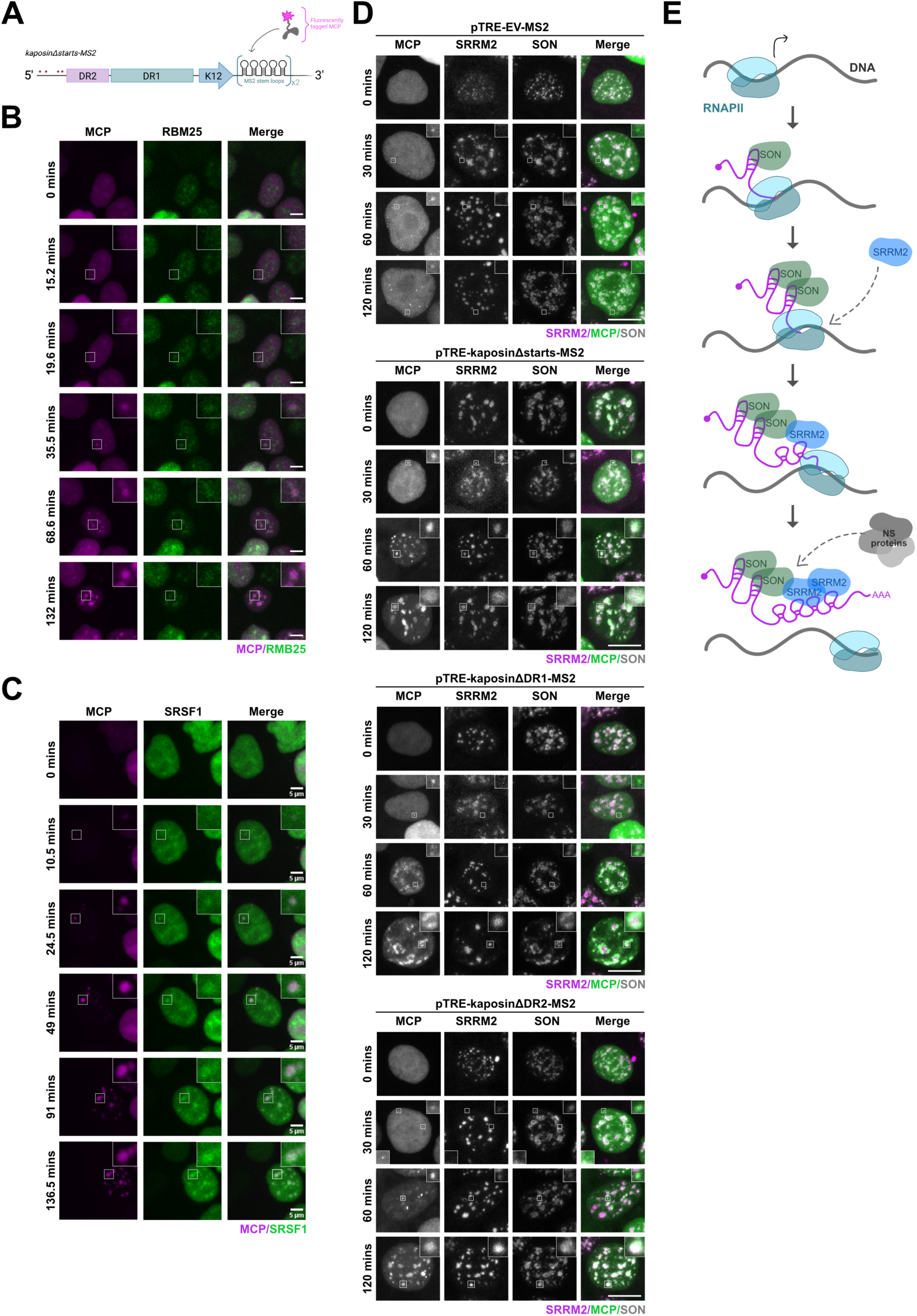
*Kaposin* transcription induces *de novo* NS assembly. **(A)** The MS2-MCP system for visualization of nascent *kaposin* RNA in real time. Table 4 for constructs. **(B-D)** 293T cells were transfected with the indicated inducible *kaposinΔstarts-MS2* construct, rtTA activator plasmid, EGFP-tagged MCP (false coloured magenta), and mCherry-tagged RBM25 (false coloured green) **(B)**/SRSF1(false coloured green) **(C)** as NS markers. For **B-C**, cells were imaged live for ∼2 h after Dox using a Nikon A1R+ confocal microscope and representative frames from Supplemental Videos 3 and 4 are shown. For **D**, cells were fixed and stained at the indicated times post induction for MCP, SRRM2 and SON. Colour threshold altered in MCP channel for visualization purposes only. **(E)** Graphical depiction of the stepwise model of *kaposin*-mediated NS seeding.

**Table 4.** Plasmid Constructs Related to **Figure 4A–D**.

| **Plasmid** | **RNA** | **KapA**<br>**ORF** | **KapB**<br>**ORF** | **KapC**<br>**ORF** | **Reference**<br>**for**<br>**construction** | **Reason for**<br>**construction** | **Figure**<br>**reference** |
| --- | --- | --- | --- | --- | --- | --- | --- |
| ΔStarts<br>(ΔS)-<br>MS2 | All canonical<br>AUG and<br>non-canonical<br>CUG start<br>codons for<br>KapA, KapB<br>and KapC<br>mutated;<br>RNA<br>contains 10 x<br>MS2 stem<br>loops at the<br>3' end | Lacks<br>start<br>AUG | Lacks<br>start<br>CUGs | Lacks<br>start<br>CUGs | This<br>manuscript | Visualization of<br>the kaposin<br>transcript using<br>MCP in live<br>imaging | Fig 4A-D |
| EV-MS2 | An empty<br>vector that<br>transcribes an<br>RNA with 10<br>x MS2 stem<br>loops | None | None | None | This<br>manuscript | Control for the<br>effect of MS2<br>stem loops on NS<br>localization | — |
| ΔSΔDR1-MS2 | All canonical<br>AUG and<br>non-canonical<br>CUG start<br>codons for<br>KapA, KapB<br>and KapC<br>mutated;<br>RNA<br>contains 10 x<br>MS2 stem<br>loops at the<br>3' end; RNA<br>lacking all<br>DR1 repeats | Lacks<br>start<br>AUG | Lacks<br>start<br>CUGs | Lacks<br>start<br>CUGs | This<br>manuscript | To determine if<br>DR1 repeats are<br>required for<br>kaposin NS<br>phenotype using<br>MCP in live<br>imaging | — |
| ΔSΔDR2-MS2 | All canonical<br>AUG and<br>non-canonical<br>CUG start<br>codons for<br>KapA, KapB<br>and KapC<br>mutated;<br>RNA<br>contains 10 x<br>MS2 stem<br>loops at the<br>3' end; RNA<br>lacking all<br>DR2 repeats | Lacks<br>start<br>AUG | Lacks<br>start<br>CUGs | Lacks<br>start<br>CUGs | This<br>manuscript | To determine if<br>DR2 repeats are<br>required for<br>kaposin NS<br>phenotype using<br>MCP in live<br>imaging | — |

**Table 5.** Plasmid Constructs Related to Supp Figure 3.

| **Plasmid** | **RNA** | **KapA**<br>**ORF** | **KapB**<br>**ORF** | **KapC**<br>**ORF** | **Reference**<br>**for**<br>**construction** | **Reason for**<br>**construction** | **Figure**<br>**reference** |
| --- | --- | --- | --- | --- | --- | --- | --- |
| BAC16<br>Forward | WT;<br>transcript<br>size | WT | WT | WT | 53 | N/A | Supp Fig<br>3 A-B |
| BAC16<br>Reverse | WT<br>sequence in<br>reverse<br>orientation<br>(3'→5') | None | None | None | This<br>manuscript | Control to<br>determine if a<br>linear sequence<br>motif in the RNA<br>is required for<br>NS phenotype | — |
| ΔS-MS2 | All canonical<br>AUG and<br>non-canonical<br>CUG start<br>codons for<br>KapA, KapB<br>and KapC<br>mutated;<br>RNA<br>contains 10 x<br>MS2 stem<br>loops at the<br>3' end | Lacks<br>start<br>AUG | Lacks<br>start<br>CUGs | Lacks<br>start<br>CUGs | This<br>manuscript | Control for MS2-MCP imaging in<br>this figure | Supp Fig<br>3 C-D |
| 4xDR2-MS2-Forward | 4 copies of<br>the 23-nucleotide<br>DR2 repeat<br>inserted<br>upstream of<br>10x MS2<br>stem loops.<br>This RNA<br>lacks kaposin<br>5'/3'<br>sequences<br>and DR1<br>repeats | None | None | None | This<br>manuscript | To determine the<br>minimal number<br>of DR2 repeats<br>for NS co-<br>localization | — |
| 4xDR1-MS2-Forward | 4 copies of the 23-nucleotide DR1 repeat inserted upstream of 10x MS2 stem loops. This RNA lacks kaposin 5'/3' sequences and DR2 repeats | None | None | None | This manuscript | To determine the minimal number of DR1 repeats for NS co-localization | — |
| 4xDR2-MS2-Reverse | 4 copies of the 23-nucleotide DR2 repeat in the reverse orientation inserted upstream of 10x MS2 stem loops. This RNA lacks kaposin 5'/3' sequences and DR1 repeats | None | None | None | This manuscript | Control to determine if a linear sequence motif in the DR2 repeat is required for NS phenotype | Supp Fig 3 E-F |
| 4xDR2-MS2-Scramble | Scrambled nucleotide sequence of 4x DR2 repeat inserted upstream of 10x MS2 stem loops. | None | None | None | This manuscript | Control for GC-rich content of the DR2 repeat | — |

| Plasmid | RNA | KapA<br>ORF | KapB<br>ORF | KapC<br>ORF | Reference<br>for<br>construction | Reason for<br>construction | Figure<br>reference |
| --- | --- | --- | --- | --- | --- | --- | --- |
| RRVrDLE2 | 4 copies of the E2 repeat from the similar RRV transcript inserted upstream of 10x MS2 stem loops. | None | None | None | This manuscript | To determine if the herpesvirus RRV encodes a GC-rich repetitive RNA that also recruits NSs | Supp Fig 3 3H |
| RRVrDLE1 | 4 copies of the E1 repeat from the similar RRV transcript inserted upstream of 10x MS2 stem loops. | None | None | None | This manuscript | To determine if the herpesvirus RRV encodes a GC-rich repetitive RNA that also recruits NSs | — |

In cells expressing *kaposinΔDR1-MS2*, MCP was more diffuse and the few foci that were observed rarely colocalized with either SON or SRRM2 at 30 min post-induction. At 1 and 2 hr post-dox, *kaposinΔDR1-MS2*-recruited MCP displayed a more punctate staining that colocalized with SON. In cells expressing *kaposinΔDR2-MS2,* MCP exhibited a mixed phenotype at early times post-induction with some small MCP puncta colocalizing with SON and SRRM2 and some not. At later times post-induction, MCP was found in large puncta that colocalized with both markers. Together these data reinforce that both DRs promote NS-colocalization and show that *kaposin* rapidly recruits NS proteins to the site of its transcription acting as the driver of *de novo* NS assembly. We propose the following model for this sequence of events: i) *kaposin* transcription is initiated by RNA polymerase II (RNAPII). ii) DR2 repeats are transcribed first and as they emerge from the RNAPII complex they are iteratively bound by SON. iii) DR1 repeats are transcribed and as they are exposed, they are iteratively bound by SRRM2 in an event required for effective *kaposin* retention in NSs. iv) Other NS proteins, such as SRSF1 and RBM25, are recruited to the complex seeded by *kaposin*, leading to complete NS compartment assembly adjacent to the site of *kaposin* transcription (Figure 4E).

### *Kaposin*-NS support viral transcription

To identify a putative function for *kaposin*-NS during KSHV infection we performed immunoprecipitation mass spec (IP-MS) on whole cell extracts of uninfected cells or reactivated iSLK-BAC16 cells using a SRRM2-specific antibody. In uninfected iSLK cells, we successfully precipitated 241 annotated NS proteins including 134/141 identified by others^11^ (Figure 5A) validating our approach. In BAC16-reactivated cells, several viral proteins were precipitated including ORF57, an RNA-binding protein that has previously been shown to localize to NSs during KSHV reactivation and enhance viral gene expression^69,70^, as well as proteins involved in viral DNA replication including K8α, ORF59, ORF6, and RTA (Figure 5B). Although their contents imply that *kaposin*-NS may be viral replication compartments, co-staining of iSLK-BAC16 cells with viral replication compartment markers, ORF6 and LANA ^71–73^, revealed *kaposin* to be adjacent to and minimally overlapping with LANA and ORF6 even at early time points (24 hours post reactivation, Supp Figure 4A-B). Similar results were obtained following co-staining for viral lncRNA *PAN* ^58,74^ (Supplemental Figure 4C). By contrast, GO analysis of statistically significantly enriched cellular proteins after SRRM2 IP-MS of infected cells revealed a strong enrichment of proteins involved in RNA transcription over that of the uninfected control condition (Figure 5C). ^22,26,24,26^ We also performed IP-MS on BAC16ΔKapB infected iSLK cells and compared the precipitated proteins to WT iSLK cells (Figure 5D).

**Figure 5.**
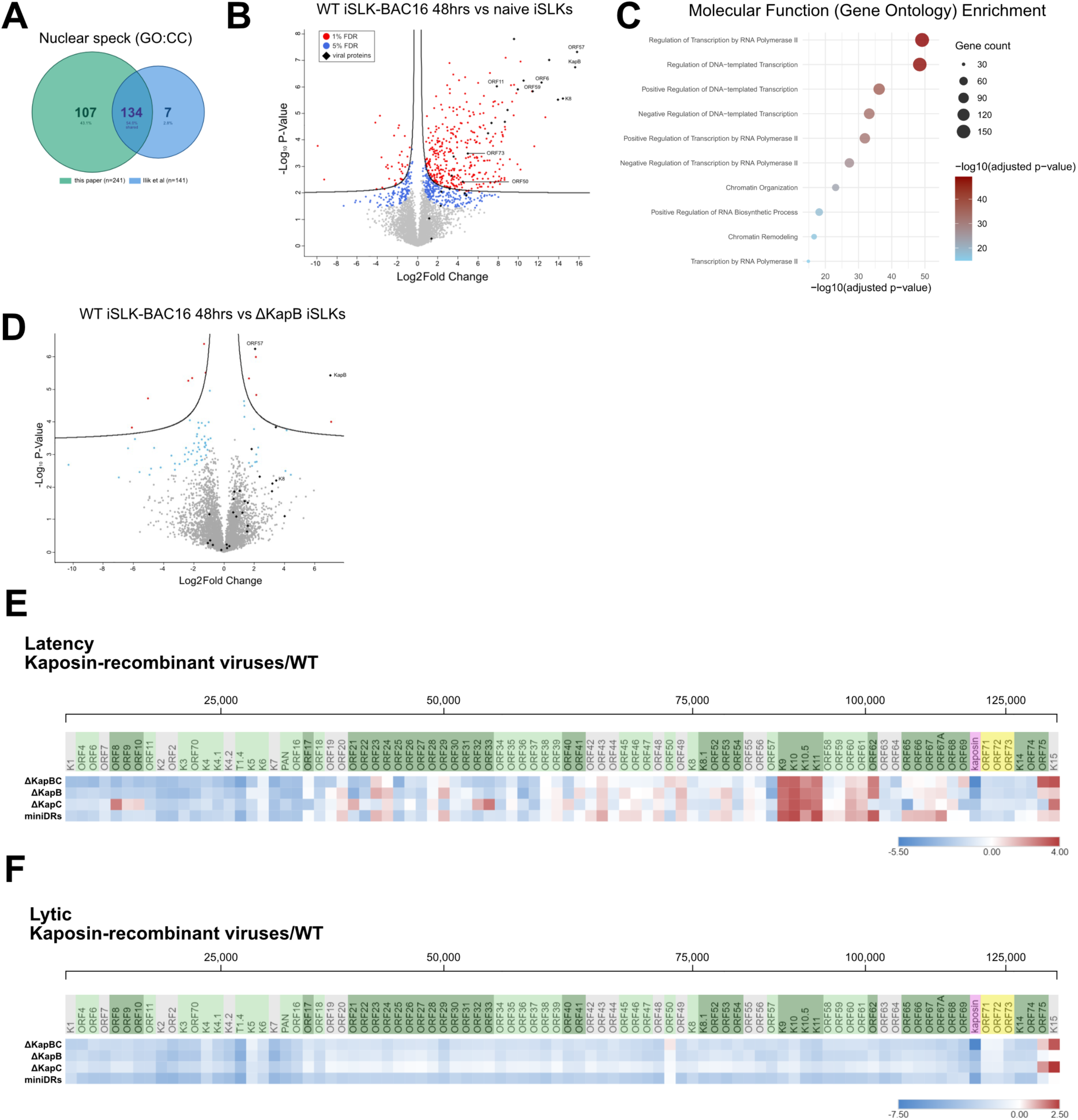
KSHV remodels NSs to optimize viral transcription. **(A-D)** Naïve or reactivated BAC16 WT/ ΔKapB iSLK cells were immunoprecipitated with anti-SRRM2 or an IgG mAb control. After validation of our NS proteome with Ilik et al. (**A**), the NS proteome was compared between WT and naïve iSLKs (**B**) or WT and ΔKapB iSLKs (**D**). Significantly enriched or depleted proteins are indicated in red (1% FDR) or blue (5% FDR); viral proteins are indicated in black. N=4 (**C**) GO analysis of proteins significantly enriched (1% FDR) in WT iSLK condition compared to naïve. Data are available via ProteomeXchange with identifier PXD080367**. (E-F)** Differentially expressed viral genes identified from RNA-seq analysis of latent (**E**) or 48 h reactivated BAC16 iSLK cells are presented as a heat map organized according to position within the viral genome, values are log2FC. Colour shading indicates temporal viral gene class, where yellow = latent, light green = early lytic, dark green = late lytic and grey = unclassed, kaposin = pink. N=5.

Several viral proteins were significantly depleted from SRRM2 precipitates including ORF57 and KapB (Figure 5D), while viral proteins involved in viral DNA replication were not significantly diminished. To test this visually, we performed IF for ORF57 and SON on naïve or infected and reactivated WT iSLK-BAC16 cells. We observed numerous instances of ORF57 colocalization with SON; however, the same colocalization was rarely observed in the matched kaposin recombinant virus conditions (Supp Figure 4D). Together, these data suggest *kaposin* is required for efficient ORF57 colocalization with NSs during KSHV infection. To determine if *kaposin* is sufficient to recruit ORF57 to NSs, we co-transfected an expression plasmid for ORF57 with increasing concentrations of the *kaposinΔstarts* plasmid. In the absence of *kaposin*, we did not observe ORF57 colocalization with SON; however, as *kaposin*-expressing plasmid concentration increased, we observed a corresponding increase in ORF57 colocalization with SON and *kaposin* (Supp Figure 4D). By independent orthogonal approaches, we reveal that *kaposin* enhances ORF57 recruitment to NSs during infection. As ORF57 supports viral gene expression, these data support that *kaposin*-NS likely function as dedicated viral transcription or RNA processing sites that are enriched in cellular transcription factors and they localize beside, but not overlapping with, viral DNA replication sites.

To determine if *kaposin*-NS seeding and remodeling is impacts viral transcription we performed RNA-sequencing to ^87^identify differentially expressed (DE) genes during WT, ΔKapBC, ΔKapB, ΔKapC, and miniDRs during latency and reactivation (Figure 5E-F). In latency, all kaposin-recombinant viruses exhibited regions of the viral genome with decreased transcription relative to WT while other regions were unchanged or increased, and these decreases did not correlate with a temporal gene expression class (latency, immediate early, early, and late) (Figure 5E). We then compared viral transcription between kaposin-recombinant viruses and WT after lytic reactivation. In this case, all *kaposin-*recombinant viruses exhibited a global decrease in gene expression relative to WT (Figure 5F), suggesting that *kaposin*-NS seeding and remodeling is required for optimal viral gene expression after reactivation.

### *Kaposin*-NS colocalization does not consistently influence mRNA export or alternative splicing of viral transcripts

NSs also promote cellular mRNA export and alternative splicing ^15,16,75–78^; therefore, we asked whether *kaposin*-mediated NS assembly similarly promotes these processes. To measure this, we reactivated latent iSLKs harbouring WT or ΔKapB, ΔKapC, and miniDRs kaposin-recombinant viruses to enter lytic replication and performed nuclear-cytoplasmic fractionation and RT-qPCR for viral mRNAs across all temporally defined KSHV gene classes: *kaposin* (latent, early), *LANA* (latent), *RTA* (immediate early), *ORF59* (early), *ORF65* (late) and *K8.1* (true late). We reasoned that if *kaposin*-NS colocalization is required for viral mRNA export we would see a decreased representation of viral transcripts in the cytoplasmic fraction during kaposin-recombinant virus reactivation relative to WT. Consistent with prior data, *kaposin* was enriched in the nuclear fraction during WT infection while *kaposin* transcripts produced by both ΔKapB and ΔKapC viruses exhibited increased cytoplasm localization (Supp Figure 5). Statistical analysis revealed no differences in the localization of *LANA*, *RTA*, *ORF65,* or *K8.1* transcripts between WT and the kaposin-recombinant viruses with the exception that *RTA* displayed increased nuclear retention during miniDRs reactivation (Supp Figure 5). These data reveal that expression of wild-type *kaposin* and NS re-positioning does not have a major consistent influence on viral mRNA export and is unlikely to be responsible for the defects associated with the kaposin recombinant viruses previously observed ^53^.To investigate whether *kaposin*-NS alter viral splicing we visualized RNA-seq read coverage across KSHV splice junctions (SJs) ^70^ during reactivation of either WT or the kaposin-recombinant viruses. We found some cases where splicing during kaposin-recombinant virus reactivation was more efficient (*K15*, SJs 51-56) and some where it was less efficient (*K10.5*, SJ 44). However, in most cases the efficiency of splicing was not noticeably altered between the WT and kaposin-recombinant virus conditions (Supp Figure 6A). We validated this *in silico* analysis using RT-PCR for *K15, K10.5,* and *K8.1* splice junctions (Supp Figure 6B-C). These data suggest that *kaposin*-mediated NS repositioning does not uniformly influence viral transcript splicing but exerts gene-specific effects. As ORF57 recruitment to NSs is impaired in the absence to WT *kaposin* (Figure 5D, Supp Figure 4D) and ORF57 NS localization contributes to its established role in viral transcript splicing ^69,70^, the loss of ORF57 from NSs may influence gene-specific changes to viral transcript splicing we observed with the kaposin-recombinant viruses.

### *Kaposin*-NS spatially regulate viral gene expression

RNA-seeded nuclear bodies have recently been appreciated to organize functional territories in the nucleus that regulate gene expression by increasing the local concentration of diffusible effector proteins in the chromatin region from which they are transcribed ^1–5^. We hypothesized that *kaposin*-mediated NS seeding acts analogously to concentrate transcription factors in physical proximity to viral genome, thereby promoting viral gene expression. To test whether *kaposin*-NS are localized near viral DNA we performed DNA-FISH in combination with RNA-FISH and IF. During latency we could detect viral genomic DNA colocalized with LANA-positive puncta (Figure 6A), validating our staining. After lytic reactivation, *kaposin* localized directly adjacent to viral DNA (Figure 6A) in SON-positive puncta (Figure 6B). To determine whether these sites displayed increased transcription, we labeled nascent RNA during lytic reactivation using 5-ethynyl uridine (EU) and observed a positive signal in *kaposin*-NS proximal regions (Figure 6C). Together these data confirm that *kaposin*-NS localize adjacent to viral DNA, positioning them appropriately to regulate viral gene expression, and that these sites correlate with increased nascent RNA abundance.

**Figure 6.**
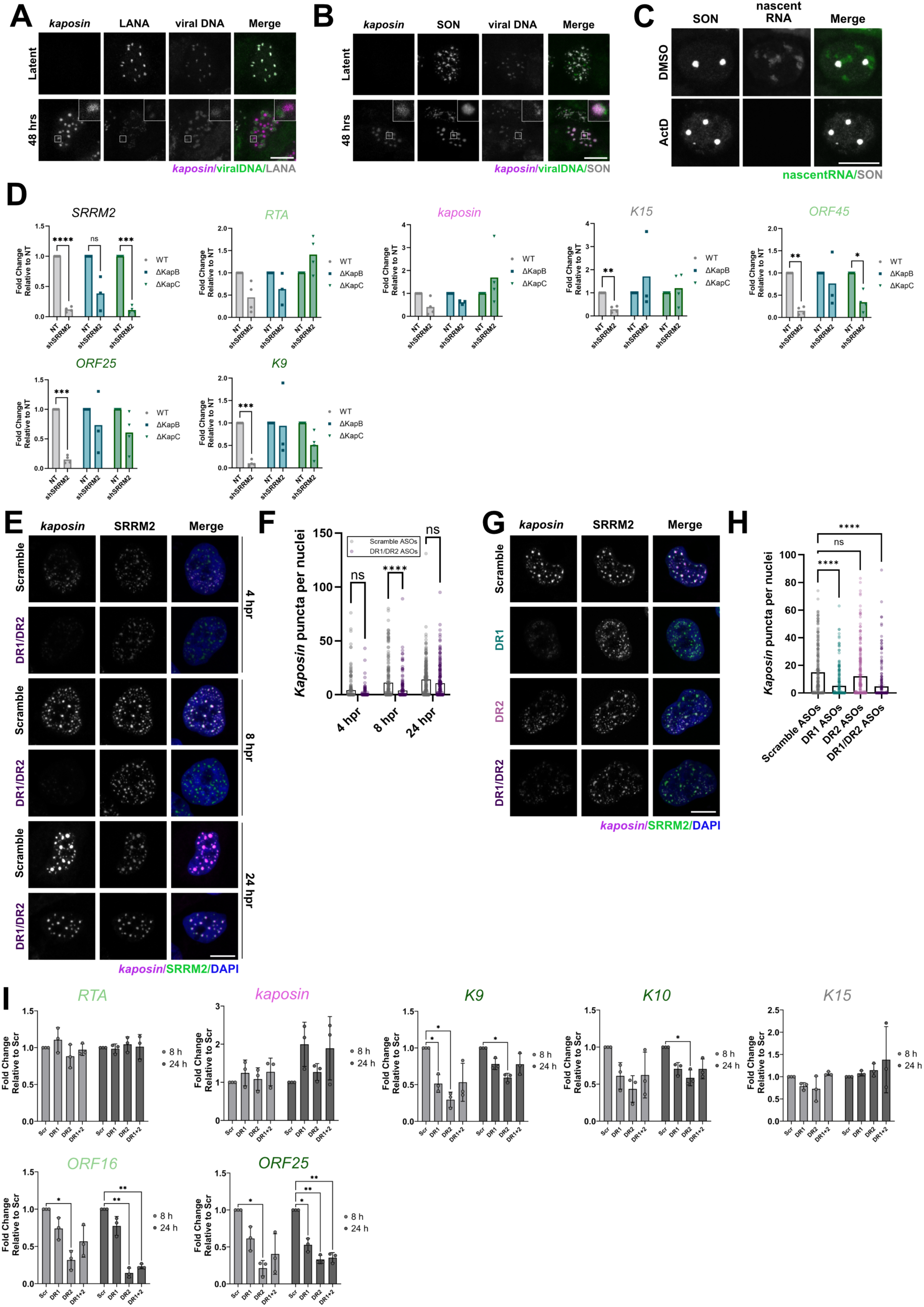
Blocking *kaposin*-NS formation impairs viral gene expression. **(A-C)** Latent or reactivated iSLK- BAC16s were stained for *kaposin*, viral DNA and either LANA (**A**) or SON (**B**) or treated with or without actinomycin D prior to 5-ethynyl uridine (EU) labeling of nascent RNA and SON **(C)**. **(D)** Latent BAC16 iSLK cells expressing SRRM2-targeting shRNAs were reactivated and RT-qPCR was performed for *SRRM2*, viral *RTA, kaposin, K15, ORF45, ORF25*, *K9* and *18S* (housekeeping). Statistics were performed using a mixed effect analysis with a Tukey’s post-hoc test. N≥3. **(E-I**) Latent BAC16 iSLK cells were reverse transfected with ASOs complementary to DR1, DR2 or a scrambled control, reactivated for indicated times, and stained for *kaposin*, SRRM2, or nuclei (**E-H**, all samples in G-H are 8 hpr). In F, H, images were quantified using CellProfiler. In parallel, RNA was isolated for RT-qPCR (**I**) as in D. Statistics were performed using a two-way repeated measures ANOVA test followed by a Dunnett’s post-hoc test (N = 3; *p* < 0.05). Temporal viral gene class: yellow = latent, light green = early lytic, dark green = late lytic and grey = unclassed, kaposin=pink.

If WT KSHV gene expression is enhanced by NS proximity, then recombinant viruses which cannot undergo *kaposin*-mediated NS repositioning, such as ΔKapB and ΔKapC (Figure 3B), should be less sensitive to NS protein depletion as they are already limited in their ability to access the NS protein pool, rendering depletion redundant. To test this hypothesis, we transduced cells latently infected with either WT, ΔKapB, or ΔKapC viruses with shRNA-expressing lentiviruses to silence SRRM2 expression. These cells were reactivated and viral gene expression was measured in comparison to a non-target (NT) shRNA control. SRRM2 knockdown during WT virus infection led to a drastic decrease in the levels of viral transcripts *K15, ORF45, ORF25,* and *K9* and a small decrease in *RTA* and *kaposin* at 48 hrs post reactivation (Figure 6D). By contrast, SRRM2 knockdown during ΔKapB and ΔKapC virus infection caused little to no change in the transcript levels of *RTA*, *kaposin*, *K15*, *ORF25*, or *K9* at 48 hours post reactivation relative to the NT controls, apart from *ORF45* transcript levels in the ΔKapC virus condition (Figure 6D). These data confirm that SRRM2 is required for optimal WT KSHV gene expression, either directly or indirectly via its role as a NS scaffolding protein, and support that a lack of NS remodeling renders viral gene expression during kaposin-recombinant virus infection less sensitive to SRRM2 depletion.

To minimize the likelihood that our observations are due to global infection state differences between WT and kaposin-recombinant viruses, we used non-degradative antisense oligonucleotides (ASOs) to sterically block the DR2 and DR1 regions of the *kaposin* transcript. After confirming ASO delivery using degradative ASOs targeting *MALAT1* (Supp Figure 7A), we transfected DR-targeting ASOs or a scrambled control DR-ASO into WT iSLK-BAC16 cells prior to reactivation and measured *kaposin* puncta formation at 4, 8, and 24 hours post reactivation. We found that steric blocking of the DRs resulted in decreased *kaposin* puncta formation across all time points, most strikingly at 8 hours (Figure 6E-F), supporting that NS protein-DR binding is required for *kaposin* condensation. Consistent with earlier findings, blocking DR1 alone had a greater effect than blocking DR2 repeats (Figure 6G-H). Similar effects were observed in our Dox-inducible *kaposin* overexpression model, reinforcing that this effect is independent of infection (Supp Figure 7C-E). Using RT-qPCR, we analyzed the transcript levels of select viral genes that were uniquely upregulated (*K9, K10, K15*) or downregulated (*ORF25, ORF16*) during the latent infection phase of kaposin-recombinant virus infection (Figure 5B). We found that steric blocking of both DRs individually or in combination significantly decreased *K9, K10, ORF16, and ORF25* transcript abundance, but did not alter the abundance of *RTA* and *K15*. These data further support *kaposin*’s role in recruitment of NS factors to boost viral transcription, recruitment specifically mediated by DR repeats. ASOs did not significantly impact transcript level of kaposin during infection or transfection models (Figure 5I, Supp Figure 7E), indicating that the mechanism of action was via steric blockade.

### *Kaposin* colocalization with NSs is required for optimal viral transcription after *de novo* **infection**

Thus far our analyses utilized the iSLK model of KSHV infection. To confirm our findings in a more physiologically relevant model, we infected human umbilical vein endothelial cells (HUVECs) ^52,53,79–81^. We reasoned that if *kaposin* was unable to reposition NSs during primary infection of HUVECs, this would impair viral transcription, explaining the decrease in viral fitness of the kaposin recombinant iSLK cell model compared to WT KSHV^53^. We first determined the infectious titer of each virus that was required to achieve equal expression of the BAC16-derived reporter, RFP, at 24 hours post infection (Figure 7A). Using samples that displayed equivalent RFP expression, we performed IF-FISH and RT-qPCR on HUVECs infected with either WT or kaposin recombinant viruses. In support of our earlier findings, *kaposin* localized adjacent to viral LANA (Figure 7B) and co-localized with SRRM2 after infection with WT but not ΔKapB or ΔKapC viruses (Figure 7C). Using RT-qPCR, we analyzed the transcript levels of the same viral genes that were upregulated (*K9, K10, K15*) or downregulated (*ORF25, ORF16*) during kaposin-recombinant virus latency (Figure 5B). At 24 hours post infection, the transcript levels of viral *RTA*, *ORF16* and *ORF25* were not significantly different between WT and kaposin recombinant viruses; however, during ΔKapB virus infection the transcript levels of *kaposin* were decreased, and during ΔKapB and ΔKapC virus infection the transcript levels of *K9* and *K10* transcripts were increased (Figure 7D). At 72 hours post infection, transcript levels of *RTA, kaposin, K9, ORF16,* and *ORF25* were significantly reduced in ΔKapB- and ΔKapC-infected HUVECs relative to WT infected cells (Figure 7E). These data provide the following important pieces of information: i) the kaposin-recombinant viruses can still enter, deliver their DNA to the nucleus, and initiate viral transcription after primary infection; ii) as early as 24 hpi, the kaposin-recombinant viral gene expression program is altered relative to WT, as evidenced by increased K9 and K10 expression, aligning with our observations during iSLK latency; iii) at 72 hpi, kaposin*-*recombinant viruses display a pronounced decrease in expression of almost all viral transcripts tested, suggesting that the lack of NS recruitment to the viral genome results in the inability of *kaposin*-recombinant viruses to sustain viral transcription after primary infection. These data suggest that *kaposin*-mediated NS repositioning plays an essential role in KSHV primary infection by promoting a favourable gene expression program. We speculate that in the absence of NS repositioning and optimal viral transcription during primary infection, latency is improperly established, as evidenced by our RNA-seq of latent iSLKs (Figure 5B) and our previous findings ^53^. This, coupled with a lack of *kaposin* RNA-driven effector protein concentration, then leads to the diminished capacity of all kaposin recombinant viruses to elicit viral transcription during reactivation (Figure 5E), an effect that was recapitulated by steric blocking of DR regions with ASOs during WT virus reactivation (Figure 6E-I). Taken together, our data support a model whereby *kaposin*-mediated RNA seeding of NS in proximity to the viral genome is required to spatially organize viral gene expression during all three stages of infection (Figure 7F).

**Figure 7.**
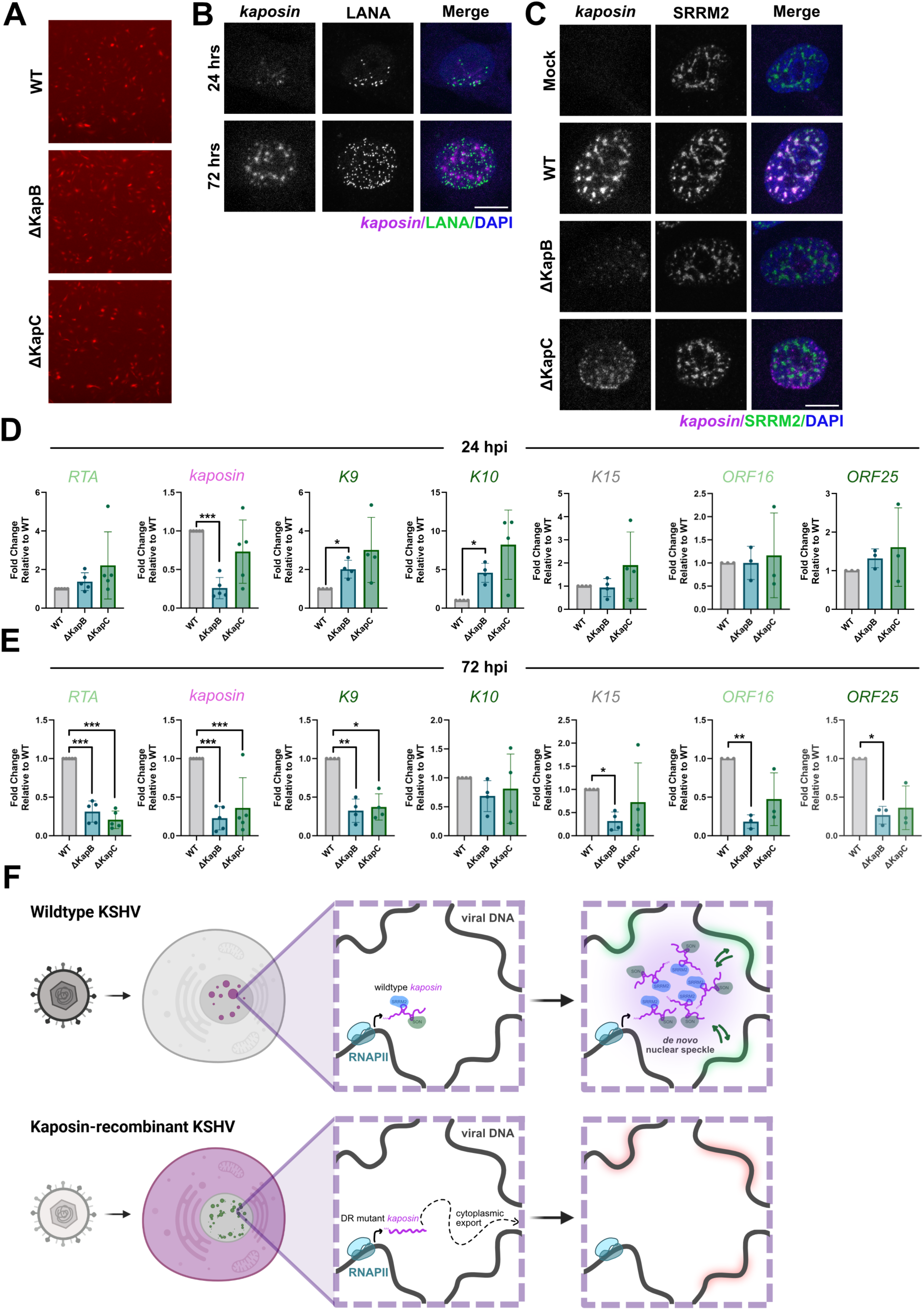
*Kaposin*-mediated NS seeding alters viral transcription after primary infection. **(A-E)** HUVECs were infected with the indicated BAC16 recombinant viruses, and RFP positivity was used to ensure equivalent infectious dose. N=5 (**A**). **(B-D)** Infected cells were stained for *kaposin* and SRRM2, and nuclei (**B,** WT only; **C** 72hpi only) or lysed for total RNA **(D)**. RT-qPCR was performed as for Figure 6. Temporal viral gene class: yellow = latent, light green = early lytic, dark green = late lytic and grey = unclassed, kaposin=pink. Statistics were performed using a repeated measures one-way ANOVA with a Tukey’s post-hoc test. N≥3; mean ± SD. **(F)** Graphical depiction of proposed model of *kaposin* function during KSHV infection.

## DISCUSSION

The three-dimensional organization of the nucleus is critical to gene expression and is heavily regulated by cellular non-coding RNAs ^1,82^. Here, we reveal a novel strategy used by a human herpesvirus to commandeer the host nucleus by employing a viral RNA to scaffold nuclear speckle (NS) assembly. We find that *kaposin* is sufficient to seed and remodel NSs, and that the *kaposin* direct repeat (DR) sequences and cellular SRRM2 are required for this effect.

Removing, altering or sterically blocking the DR sequences leads to a loss of *kaposin-*NS colocalization during KSHV infection and altered transcription of viral genes. These data indicate that KSHV uses *kaposin* as a viral architectural (arc)RNA to effectively remodel the infected cell nucleus for optimal gene expression.

### A new role for the *kaposin* transcript in KSHV infection

Despite the historical emphasis placed on Kap protein function ^47,48,50–52^, *kaposin* is absent from polysomes during latent iSLK infection ^26^ and in most KS tumours sequenced the *kaposin* RNA is predicted to be non-coding ^45^. The conservation of the nucleotide sequence of *kaposin* over the Kap ORFs^45^ suggests *kaposin* encodes an RNA-based function which supersedes that of its cognate proteins. We now show that the majority of the *kaposin* RNA is retained in the nucleus where it colocalizes with NSs. In a minority of reactivated cells, we observe cytoplasmic *kaposin* staining alongside a strong nuclear *kaposin* signal. We speculate that abundant *kaposin* transcription overwhelms the nuclear pool of available SRRM2 and, as a result, *kaposin* is no longer retained in NSs, allowing it to be exported and subsequently translated. Kap proteins are likely only expressed by a subset of cells undergoing robust reactivation or transfection conditions where enough RNA has accumulated to permit NS escape. This observation suggests that Kap proteins may function in late lytic replication, not latency, as previously assumed.

Several herpesviruses encode GC-rich and highly repetitive RNAs from similar genomic positions ^83–87^. Here, we find that three repeat units from the DR2-like region of RRV are sufficient to promote NS localization while work from others has shown that some of these RNAs also exhibit punctate nuclear staining during infection and, in the case of Epstein-Barr Virus (EBV), colocalize with SRRM2 and are adjacent to viral replication compartments ^21,88^. These observations suggest that scaffolding of nuclear bodies by a viral arcRNA may be a universal characteristic of herpesvirus infection.

### *Kaposin* is a viral arcRNA that seeds nuclear speckles

Despite significant study, a cellular arcRNA capable of seeding NSs has yet to be identified. Using a combination of fixed cell and real time imaging, we now show that the viral RNA *kaposin* is sufficient to drive formation of a NSs *de novo* and remodel their composition during infection. We propose that the DR1 repeat of *kaposin* functions as the primary NS retention signal that directly or indirectly binds to the inner NS scaffolding protein, SRRM2, and the DR2 repeat encodes a secondary signal responsible for binding and recruiting SON. SRRM2 binding has been reported to be non-specific ^12^; however, we speculate that the key to the strong recruitment by *kaposin* is the number of DR1 repeats. 27 repeats of this sequence are found in WT *kaposin*, while the miniDRs recombinant virus/construct contains only five copies of DR1 and retains partial SRRM2 recruitment capability. These data suggest that each DR1 unit forms a structure that when iteratively repeated is sufficient for strong SRRM2 binding specificity.

### *Kaposin*-NSs optimize viral transcription

Systematic mapping of genomic-NS contacts reveals that NS positioning correlates tightly with gene expression ^75,76^. Transcriptional boosting is thought to occur via decreased diffusion distance of NS-supplied RNA processing enzymes. If KSHV co-opts NS function to enhance viral transcription, NSs should be proximal to viral DNA and should contain cellular and viral proteins known to enhance transcription. We show that both events occur. NSs are found beside viral DNA during infection and kaposin-NSs are enriched in cellular and viral (ORF57) proteins that promote transcription. Using RNA sequencing to compare between KSHV viruses that cannot remodel NS with those that can, we show the lack of *kaposin*-NS colocalization dysregulates viral transcription following a pattern based on genomic position in latency and early primary infection. During lytic reactivation and late primary infection, the absence of *kaposin*-NSs corresponds to a global decrease in viral gene expression. To ensure these data were not influenced by global infection state, we used non-degradative ASOs to sterically block DR1 and DR2 regions during viral reactivation and showed that blocking DR1 alone or both DRs prevented formation of *kaposin*-NSs and decreased viral transcription. Together, these data show that *kaposin*-NSs congregate viral and cellular transcription machinery beside the viral DNA to optimize viral gene expression.

### Limitations

We acknowledge the following limitations of our study. i) Experiments using real-time imaging of nascent *kaposin* after induction show *kaposin* recruits NS proteins to its transcription site, indicating *kaposin* is the driver or ‘seed’ for *kaposin*-NSs, and not the passenger. However, we cannot distinguish whether *kaposin* is the driver or a passenger in *kaposin*-NSs in fixed cell models. ii) Despite extensive mutagenesis of *kaposin*, which identified DR1 as an essential and DR2 as a supporting signal for *kaposin*-NS co-localization, we have not demonstrated direct binding of NS proteins to these regions. iii) We were surprised to observe that *kaposin* constructs devoid of known start codons (*kaposinΔstarts*) produce unknown translation products after overexpression. GC-rich repeat RNAs often translate non-canonical products using alternative mechanisms^63,64^. Although these products cannot be translated in our inducible *kaposin* system, where NS proteins are recruited to the nascent transcript as early as 19 minutes after induction, we cannot discount a minor contribution of the Kap polypeptides to NS remodeling at later transfection times or during infection. iv) Our data provide compelling evidence for *kaposin*-mediated recruitment of NS components adjacent to the viral genome, supporting a model where NS recruitment is the required event to boost viral transcription. However, our data does not yet fully exclude alternative or parallel mechanisms by which *kaposin* may alter transcription factors or chromatin modifications to boost viral transcription.

## Supporting information

Supplemental Data table related to Figure 5

## Acknowledgements

The authors would like to dedicate this work to Dr. Robert Sadler, who was the first person to characterize the complicated KSHV *kaposin* transcript. The authors sincerely thank the members of the Corcoran lab for helpful discussions and Drs. Craig McCormick and Roy Duncan (Dalhousie University) for providing constructive feedback on the manuscript. We thank Dr. Anne Vaahtokari, Mr. Dylan Greening, the Charbonneau Institute Microscopy Facility, and the Snyder Institute Live Cell Imaging Laboratory for microscopy support and the Cumming School of Medicine, Centre for Health Genomics and Informatics, for RNA sequencing support.

Plasmids kindly gifted to us through Addgene are detailed in Table 3. MK was supported by a Canadian Institutes for Health Research (CIHR CGS-D) Canada Graduate Scholarship and a Cumming School of Medicine (CSM) doctoral training award. RPM was supported by a Snyder Institute Beverley Phillips Doctoral training award and a CIHR CGS-D award. MPBM was supported by a CSM doctoral training award, a Snyder Institute Beverley Phillips doctoral training award and an Alberta graduate scholarship. Operating funds to support this work derive from a Canadian Institutes for Health Research Project Grant PJT-183595 awarded to JAC. Graphics were created using BioRender or Affinity Designer.

## Author Contributions

Mariel Kleer: Conceptualization, Experimentation, Data Analysis, Paper Writing, Paper Editing Rory P. Mulloy: Experimentation, Data Analysis, Paper Editing

Maxwell P. Bui-Marinos: Experimentation, Data Analysis, Paper Editing Michael Johnston: Data Analysis, Paper Editing

Morgan F. Khan: Experimentation, Data Analysis, Paper Editing David C. Schriemer: Supervision, Paper Editing

Jennifer A. Corcoran: Conceptualization, Experimentation, Supervision, Funding Acquisition, Project Administration, Paper Writing, Paper Editing

## METHODS

### Cell Culture

All cells were grown at 37°C with 5% CO_2_ and atmospheric O_2_. HEK293T and HeLa cells (ATCC) were cultured in Dulbecco modified Eagle medium (DMEM; ThermoFisher) supplemented with 100 U/mL penicillin, 100 μg/mL streptomycin, 2 mM L-glutamine (ThermoFisher), and 10% fetal bovine serum (FBS; ThermoFisher). Naïve iSLK-TREx-RTA cells ^55^ were cultured likewise with the addition of 1 μg/mL puromycin and 250 μg/mL Geneticin (ThermoFisher). Naïve iSLK-TREx-RTA cells stably infected with KSHV BAC16 (BAC16-iSLK-TREx-RTA) were cultured as naïve iSLK-TREx-RTA cells with the addition of 1200 μg/mL hygromycin B (ThermoFisher). BCBL1-TREx-RTA cells were cultured in RPMI 1640 (ThermoFisher) supplemented with 55 μM β-mercaptoethanol, 100 U/mL penicillin, 100 µg/mL streptomycin and 10% FBS. For immunofluorescence BCBL1-TREx-RTA cells were seeded into glass coverslips coated with 50 μg/mL poly-D-lysine (Sigma-Aldrich). Human umbilical vein endothelial cells (HUVECs; Lonza) were cultured in endothelial cell growth medium (EGM-2; Lonza). HUVECs were seeded onto gelatin (0.1% [wt/vol] in PBS)-coated tissue culture plates or glass coverslips.

### Production of KSHV

BAC16-iSLK-TREx-RTA cells were reactivated using 1 μg/mL doxycycline (Dox; Sigma-Aldrich) and 1 mM sodium butyrate (NaB; Sigma-Aldrich) in antibiotic-free 10% FBS–DMEM. At 72h post reactivation, supernatants were collected, clarified at 1200 × g to remove cellular debris, and virus-containing supernatant was stored at −80°C prior to use. Because an RFP reporter gene is encoded by the BAC16 genome under a mammalian promoter, the infectious titer for both WT and kaposin-recombinant KSHV was estimated using the number of RFP-positive cells 24 h post infection of 293T cells.

### BAC16 mutagenesis and stable iSLK cell line generation

To create the miniDRs recombinant virus Lambda Red recombination was carried out in the BAC16 background ^54^ supplied in the GS1783 strain of *Escherichia coli* (a generous gift from Dr. Jae Jung, Lerner Research Institute, Cleveland Clinic) as extensively detailed in^53^. The stable iSLK cell line harbouring the miniDRs genome was likewise created as previously described ^53^.

### Primary infection

Subconfluent (50 to 70%) 293T or HUVEC cells were incubated with viral inoculum diluted in antibiotic and serum-free DMEM containing 8 μg/mL hexadimethrine bromide (Polybrene; Sigma-Aldrich). Plates were centrifuged at 800 × g for 1.5 h at room temperature and the inoculum was removed and replaced with either antibiotic-free 10% FBS–DMEM (293T) or full EGM-2 (HUVECs). Cells were then returned to the 37°C incubator.

### IF-FISH

IF-FISH was performed according to manufacturer’s protocol (Stellaris IF-FISH protocol). Briefly, cells were fixed in 4% paraformaldehyde (Sigma-Aldrich) for 10 min then permeabilized with 0.1% Triton X-100 (Sigma-Aldrich) in PBS for 10 min. Primary antibodies (Table 6) were diluted in PBS and incubated with cells for 1 h at RT, after which cells were washed and corresponding secondary antibodies (Table 1) were likewise diluted in PBS and incubated with cells for 1 h at RT. After IF, cells were fixed in 4% paraformaldehyde for 10 min, washed with PBS, and incubated in Wash Buffer A (LGC Biosearch) for 5 min. Stellaris probes (Table 7) were diluted in Hybridization Buffer (LGC Biosearch) to a concentration of 125 nM and cells were incubated for hybridization at 37°C overnight. The following day, cells were washed sequentially with Wash Buffer A for 30 min, Wash Buffer A containing 5 ng/mL DAPI (Invitrogen) for 30 min, and Wash Buffer B (LGC Biosearch) for 5 min or, for non DAPI stained samples, just Wash Buffer A and B. Samples were mounted with Prolong Gold AntiFade mounting media (Thermo Fisher) and imaged on a Zeiss LSM 880 laser scanning confocal microscope using the 63X 1.4 NA Oil Plan-Apo objective and the following lasers (excitation wavelengths): Diode (405nm), Argon (488nm), and Helium/Neon (561nm and 633nm) with emission filter ranges of 410-513nm, 489-605nm, 570-659nm and 638-755nm, respectively.

**Table 6:** Antibodies:

| Antibody | Species | Vendor/Catalog | Application | Dilution |
| --- | --- | --- | --- | --- |
| SRRM2 | Mouse | Sigma Aldrich, S4045 | IP | 1:200 |
| IgG isotype control | Mouse | CST | IP | 1:200 |
| SRRM2 | Mouse | Abcam, ab11826 | IF | 1:200 |
| SON | Rabbit | Abcam, ab121759 | IF | 1:200 |
| PML | Mouse | Santa Cruz, sc-966 | IF | 1:200 |
| Coilin | Mouse | Abcam, ab87913 | IF | 1:1000 |
| PRPF8 | Rabbit | Abcam, ab79237 | IF | 1:200 |
| LANA | Rabbit | A generous gift from Dr. Craig McCormick | IF | 1:1000 |
| ORF57 | Mouse | Santa Cruz, sc-135746 | IF | 1:1000 |
| ORF6 | Rabbit | A generous gift from Dr. John Karijovich | IF | 1:1000 |
| Tubulin | Rabbit | CST 2128S | immunoblotting | 1:1000 |
| Lamin A/C | Rabbit | CST 2032S | immunoblotting | 1:1000 |
| Kaposin B/C | Rabbit | A generous gift from Dr. Craig McCormick | immunoblotting | 1:1000 |
| Chicken anti Rabbit Alexa Fluor™ 488 | Chicken | ThermoFisher, A21441 | IF | 1:1000 |
| Goat anti Mouse Alexa Fluor™ 488 | Goat | ThermoFisher, A11029 | IF | 1:1000 |
| Donkey anti Rabbit Alexa Fluor™ 405 | Donkey | Abcam, ab175651 | IF | 1:200 |

**Table 7:**
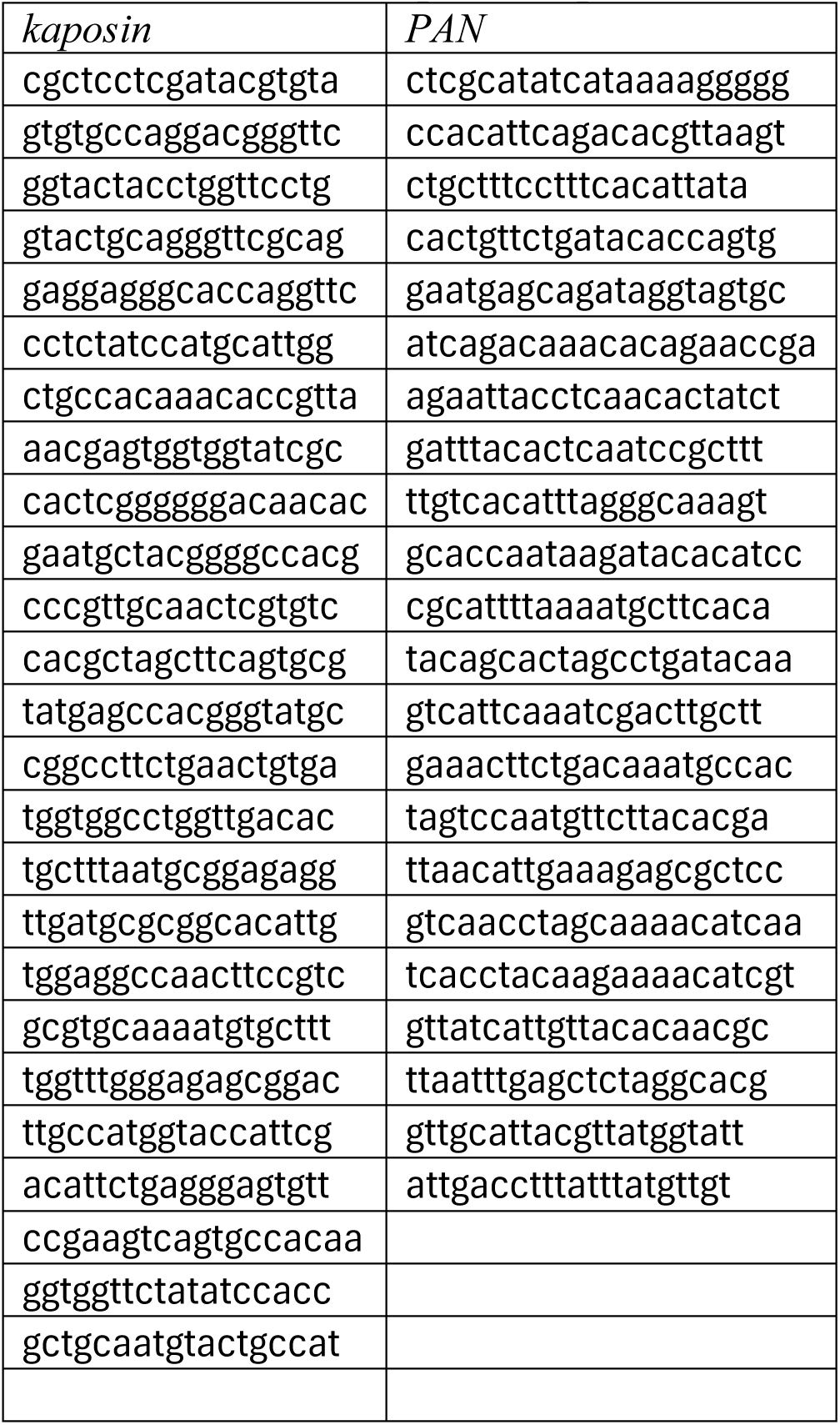
Custom FISH probe sequences.

Maximal intensity projections (MIPs) with dimensions of 1912 x 1912 pixels are presented unless otherwise specified. Color thresholding was adjusted equally within experiments except for Figure 3B (WT, kaposin channel), Figure 4B (MCP channel), and Supplemental Figure 3D (MCP channel) which were altered purely for visualization purposes due to heterogeneity of expression. Altered images were never used for quantification. Scale bars = 10 uM.

### Live cell imaging

Cells were seeded and transfected as described in the associated figure captions. 24 hours post transfection, the media was replaced with phenol-free DMEM (Gibco), supplemented with 100 U/mL penicillin, 100 μg/mL streptomycin, 2 mM L-glutamine (ThermoFisher), and 10% fetal bovine serum (FBS; ThermoFisher). Cells were transported onto Nikon A1R confocal microscope equipped with live-cell imaging chamber preset to 37°C with 5% CO_2_ and atmospheric O_2_. Cells were left to acclimate for 30 minutes. Prior to imaging, cells were treated with 1 µg/mL doxycycline (Dox; Sigma-Aldrich). Cells were imaged using 20X air objective at the indicated intervals. Time-lapse videos of maximum intensity projections were produced using ImageJ. Color thresholding was adjusted equally within experiments. Scale bar = 5 uM.

### BACmid DNA extraction and KSHV DNA probe synthesis

BACmid DNA was isolated from GS1783 E. coli cultures of 500 mL of LB containing 30 μg/mL chloramphenicol (Sigma-Aldrich) grown overnight at 30°C. Bacteria were pelleted at 4,000 × g for 10 min at 4°C and DNA was extracted using the Qiagen large construct kit according to manufacturer’s instructions. Probes for the detection of KSHV BACmid DNA were synthesized using the BioNick™ DNA labeling system (ThermoFisher) according to manufacturer’s protocol with the following modifications. Briefly, the reaction was scaled up to 3 µg and allowed to proceed for 2 h, instead of 1, prior to the addition of stop buffer. Probes were then purified using repeated ethanol precipitation, followed by column purification and resuspension in 30 µL of TE buffer and stored at −20°C prior to use.

### DNA IF-FISH

Cells were fixed in 4% paraformaldehyde (Sigma-Aldrich) for 10 min then permeabilized with 0.1% Triton X-100 (Sigma-Aldrich) in PBS for 10 min. Cells were washed with PBS, dehydrated using a series of ethanol concentrations (70, 80, 90, and 100%), and air dried for 10 min at RT prior to incubation with hybridization solution (CytoCell) containing 3 µL reconstituted BACmid DNA probes (generated as above) and Stellaris RNA probes to a final concentration of 125 nM. The coverslips/hybridization mixture was sealed cell side down to the slide using rubber cement and denatured at 83°C for 8 min prior to incubation at 37°C overnight. The following day coverslips were removed from the slide, washed 2X for 15 min at RT with 70% (vol/vol) formamide/10 mM Tris-HCl [pH 7.2] and 3X for 5 min at RT with 0.1 M Tris-HCl [pH 7.2]/0.15 M NaCl containing 0.05% (vol/vol) Tween 20. Cells were washed with PBS and Avidin, NeutrAvidin™, Oregon Green™ 488 conjugate (ThermoFisher) was diluted in 2% BSA in PBS and incubated for 1 h at RT. Cells were washed with PBS and primary antibodies (Table 1) were diluted in PBS and incubated for 1 h at RT. Cells were washed with PBS and corresponding secondary antibodies (Table 6) were likewise diluted in PBS and incubated for 1 h at RT. Samples were mounted with Prolong Gold AntiFade mounting media (ThermoFisher) and imaged on a Zeiss LSM 880 laser scanning confocal microscope using the 63X 1.4 NA Oil Plan-Apo objective and the following lasers (excitation wavelengths): Diode (405nm), Argon (488nm), and Helium/Neon (561nm and 633nm) with emission filter ranges of 410-513nm, 489-605nm, 570-659nm and 638-755nm, respectively. Maximal intensity projections (MIPs) with dimensions of 1912 x 1912 pixels are presented unless otherwise specified.

### Nascent RNA imaging

Nascent RNA was visualized using a Click-iT™ RNA Alexa Fluor™ 488 Imaging Kit (Invitrogen, C10329) according to standard protocol. Briefly, cells were treated with 5-ethynyl uridine (EU) for 1 hour prior to fixation. Cells were then permeabilized, washed, and Click-iT® reaction cocktail was added and incubated for 30 minutes. Cells were washed once with Click-iT® reaction rinse buffer and IF labeling was performed.

### Nuclear cytoplasmic fractionation and DNase treatment

Nuclear cytoplasmic fractionation was conducted as in ^74^. Briefly, cells were lysed in 300 uL Nuclear Extraction Buffer (NEB; (20 mM HEPES [pH 7.2], 50 mM NaCl, 3 mM MgCl_2_, 300 mM sucrose, and 0.5% NP-40) for 15 min on ice and collected by scraping. Lysates were clarified by centrifugation at 10 000 x g for 10 min at 4°C and the supernatant containing the cytoplasmic fraction was removed and set aside. The nuclear pellet was washed in 500 uL ice-cold PBS and spun down at 10 000 x g for 10 min at 4°C. The PBS wash was removed, and the pellet was resuspended in 300 uL NEB. Both cytoplasmic and nuclear fractions were disrupted using a probe sonicator for 10 s 2X each. 900 uL of TRIzol (Thermo Fisher) was added to each sample and RNA was extracted according to manufacturer’s instructions. Genomic DNA was removed using the Turbo DNA-free kit (Thermo Fisher) and the resulting RNA was stored at −80°C until further use.

### Nuclear/cytoplasmic fractionation and TaqMan PCR analysis

10-cm dishes of transfected 293Ts or 48 hour-reactivated WT BAC16/naïve iSLKs were washed in of PBS and lysed in 1ml of ice-cold nuclei EZ lysis buffer from the Nuclei EZ Prep nuclei isolation kit (Sigma Aldrich Cat# NUC-101) supplemented with protease inhibitor cocktail (Sigma Aldrich) and 60 U/mL of SUPERase In™ RNase Inhibitor (ThermoFisher). Cells were scraped, collected, and set on ice for 5 min. Nuclei were collected by centrifugation at 500xg at 4C for 5 minutes. 600ul of the clarified supernatant was reserved as the cytoplasmic fraction; the remainder was discarded so as not to contaminate fractions. Pelleted nuclei were washed twice in 2ml of ice-cold nuclei EZ lysis buffer. Washed nuclear pellets were lysed in 600ul of Pull Down (PD) buffer (20mM Tris–HCl pH 8.0, 200mM NaCl, 2.5mM MgCl2, 0.05% Triton X-100 containing protease inhibitors and 60 U/mL of SUPERasel In™ RNase Inhibitor) ^89^. From each 600ul fraction, 500 ul was mixed with 500ul of TRIzol LS (ThermoFisher) for RNA extraction and 100ul was mixed with 4x Laemelli buffer for immunoblotting (described below).

RNA was extracted by TRIzol protocol, and contaminating DNA removed using TURBO DNase (ThermoFisher Cat# AM2238), according to manufacturer’s specifications. RNA quality was determined using NanoDrop OneC (ThermoFisher) and concentration was determined using Qubit RNA Broad Range assay (ThermoFisher Cat# Q10210); 1000 ng of RNA was reverse transcribed using MaximaH Minus Master Mix (ThermoFisher Cat# M1661) in a 10 μL reaction (100 ng RNA/μL). cDNA was diluted 1:50 (293T; represents 2 ng RNA/μL) or 1:10 (iSLK, represents 10 ng RNA/μL) in water, of which 2.5 μL was used in qPCR, meaning each qPCR reaction is estimated to represent 5 or 25 ng of RNA, respectively.

The *kaposin* TaqMan probe was custom designed by providing a 164-nucleotide sequence from the KapA region of kaposin (TGTTTGTGGCAGTTCATGTCCCGGATGTGTTACTAAATGGGTGGCGCTGGAGGCTTGGGGCGATACCACCACTCGTTTGTCTGTTGGCGATTAGTGTTGTCCCCCCGAGTGGCCAGCGTGGCCCCGTAGCATTCAGGACACGAGTTGCAACGGGCGCGCACTGA; Assay ID AP7D4GA) to Applied Biosystems Custom TaqMan Gene Expression service (ThermoFisher). Probes were end-labeled with a dye label (FAM) on the 5’ end and a minor groove binder (MGB) and nonfluorescent quencher (NFQ) on the 3’ end; two unlabeled primers were designed to flank the probe-binding site, allowing for greater accuracy and specificity in detecting *kaposin* transcripts. qPCR reactions were prepared in TaqMan Universal PCR Master Mix (ThermoFisher, Cat# 4304437) as per manufacturer’s instructions, using 2.5 μL of diluted cDNA per reaction. Cq values were related against a standard curve of known pcDNA-kaposin-ΔStarts plasmid copies, equating to the number of *kaposin* transcript copy numbers. Resulting reaction transcript copy number (TCN) were divided by the amount of RNA represented in the qPCR reaction to infer the concentration of *kaposin* transcripts within the given compartment RNA pool.

### Transfection

293T cells seeded to 70% confluency in antibiotic-free 10% FBS–DMEM were washed once with PBS, and medium was replaced with 800 uL of serum-free, antibiotic-free DMEM per well prior to transfection. Cells were transfected with plasmids of interest (Table 8) using polyethylenimine (PEI, Polysciences). At 4 h post-transfection, the medium was replaced with antibiotic-free 10% FBS–DMEM.

**Table 8:** Plasmids.

| Plasmid | Use | Source |
| --- | --- | --- |
| pcDNA3.1(+) | Empty Vector (EV) control | Invitrogen V79020 |
| pcDNA-kaposin | Overexpression | BioBasic custom gene synthesis |
| pcDNA-kaposin-reverse | Overexpression | Cloned into pcDNA using NheI and BamHI |
| pcDNA-kaposin-ΔStarts | Overexpression | BioBasic custom gene synthesis |
| pcDNA-kaposin-altB | Overexpression | BioBasic custom gene synthesis |
| pcDNA-kaposin-ΔStarts-ΔDR1 | Overexpression | BioBasic custom gene synthesis |
| pcDNA-kaposin-ΔStarts-ΔDR2 | Overexpression | BioBasic custom gene synthesis |
| pcDNA-kaposin-ΔStarts-DRswap | Overexpression | BioBasic custom gene synthesis |
| pcDNA-kaposin-miniDR | Overexpression | BioBasic custom gene synthesis |
| pcDNA-kaposin-miniDR-swap | Overexpression | BioBasic custom gene synthesis |
| pcDNA-kaposin-5X-DR1 | Overexpression | BioBasic custom gene synthesis |
| pcDNA-kaposin-3X-DR2 | Overexpression | BioBasic custom gene synthesis |
| pcDNA-rtTA | Overexpression | Cloned into pcDNA using NheI and NotI |
| MS2_GFP | Cloning | MS2_GFP was a gift from Daniel Larson (Addgene plasmid # 61764) |
| MCP-NLS-EGFP | Overexpression | Cloned from MS2_GFP using BamHI and NotI |
| pEGFP-SF2 | Cloning | pEGFP-SF2 was a gift from Tom Misteli (Addgene plasmid # 17990) |
| mCherry-SRSF1 | Overexpression | Cloned from pEGFP-SF2 using HindIII and BamHI |
| mCherry-RBM25 | Overexpression | Cloned from cDNA using NheI and SalI |
| pTRE-SOX | Cloning | pTRE-SOX was a gift from Dr. Eric Pringle |
| pcDNA-puro-TagBFP-β-ACTIN-3'UTR-24xMS2 | Cloning | pcDNA-puro-TagBFP-β-ACTIN-3'UTR-24xMS2 was a gift from Weirui Ma (Addgene plasmid # 20516) |
| pTRE-EV-MS2 | Cloning/Overexpression | Cloned from pTRE-SOX and pcDNA-puro-TagBFP-β-ACTIN-3'UTR-24xMS2 |
| pTRE-kaposin-ΔStarts-MS2 | Overexpression | Cloned into pTRE-EV-MS2 from pcDNA-kaposin-ΔStarts using NheI and EcoRI |
| pTRE-kaposin-ΔDR1-MS2 | Overexpression | Cloned into pTRE-EV-MS2 from pcDNA-kaposin-ΔStarts-ΔDR1 using NheI and EcoRI |
| pTRE-kaposin-ΔDR2-MS2 | Overexpression | Cloned into pTRE-EV-MS2 from pcDNA-kaposin-ΔStarts-ΔDR2 using NheI and EcoRI |
| pTRE-4X-DR2-MS2 | Overexpression | Cloned into pTRE-EV-MS2 from pTRE-EV-MS2 used annealed oligos with NheI and EcoRI sticky ends |
| pTRE-4X-DR1-MS2 | Overexpression | Cloned into pTRE-EV-MS2 from pTRE-EV-MS2 used annealed oligos with NheI and EcoRI sticky ends |
| pTRE-4X-DR2-reverse-MS2 | Overexpression | Cloned into pTRE-EV-MS2 from pTRE-EV-MS2 used annealed oligos with NheI and EcoRI sticky ends |
| pTRE-4X-DR2-scramble-MS2 | Overexpression | Cloned into pTRE-EV-MS2 from pTRE-EV-MS2 used annealed oligos with NheI and EcoRI sticky ends |
| pTRE-RRV-3X-rDL-E2-MS2 | Overexpression | Cloned into pTRE-EV-MS2 from pTRE-EV-MS2 used annealed oligos with NheI and EcoRI sticky ends |
| pTRE-RRV-5X-rDL-E1-MS2 | Overexpression | Cloned into pTRE-EV-MS2 from pTRE-EV-MS2 used annealed oligos with NheI and EcoRI sticky ends |
| pcDNA-ORF57 | Overexpression | BioBasic custom gene synthesis |
| psPAX2 | Lentivirus production | psPAX2 was a gift from Didier Trono (Addgene plasmid # 12260) |
| pMD2.G | Lentivirus production | pMD2.G was a gift from Didier Trono (Addgene plasmid # 12259) |
| pLKO.1-blast | Cloning | pLKO.1-blast was a gift from Keith Mostov (Addgene plasmid # 26655) |
| pLKO-NT | shRNA expression | A gift from Dr. Craig McCormick |
| pLKO-SRRM2-sh1 | shRNA expression | Cloned into pLKO.1-blast using AgeI and EcoRI |
| pLKO-SRRM2-sh2 | shRNA expression | Cloned into pLKO.1-blast using AgeI and EcoRI |
| pCDH-hMALAT1 | MALAT1 standard curve | pCDH-hMALAT1 was a gift from Jianjun Zhao (Addgene plasmid # 118580) |

### Immunoblotting

Following cytoplasmic/nuclear fractionation, cell lysate was stored at lysed in 2x Laemmli −20°C. Protein concentration was determined using DC Protein Assay kit (BioRad). 6 µg of nuclear protein fraction or 3 µg of cytoplasmic protein fraction was resolved by SDS-PAGE using a TGX Stain-Free acrylamide gel (BioRad). Following transfer, PVDF membranes were blocked with 5% BSA (Thermo-Fisher) in TBST (Tris-buffered saline solution, 0.1% Tween-20). Primary antibodies were diluted in 2.5% BSA in TBST, and secondary antibodies were diluted in 5% skim milk in TBST. Antibody dilutions can be found in Table 6. Total protein and chemiluminescent images were acquired on a ChemiDoc MP Imaging System (BioRad).

### Antisense Oligonucleotide (ASO) experiments

Antisense oligonucleotides (ASOs) modified with 2’-O-methoxy-ethyl (2’-MOE) were custom designed by Integrated DNA technologies (IDT) against *kaposin* direct repeat regions. The 2’-MOE modification increases resistance to nuclease degradation, reduces toxicity, and increases affinity for binding to complimentary RNA. A control gapmer ASO against the cellular nuclear speckle lncRNA, *MALAT1*, was used as a positive control for ASO nuclear delivery (Table 13). 293Ts, WT BAC16 iSLKs, or naïve iSLK cells were reverse transfected using lipofectamine 2000 (Invitrogen) and the indicated concentration of ASO (for MALAT1: 10nM, 3nM, 1nM, 0.3nM, 0.1nM; for DR1/DR2/DRCtrl: 10nM; for DR1+DR2:10nM total [5nm each DR1 and DR2 ASO]). For 293Ts, coverslips were pre-treated with 10ug/ml fibronectin for 30 minutes at 37C. To test *kaposin* blockade, 293T cells were subsequently transfected with 900ng pTre-kaposin delStarts/empty vector control and 100ng of rTA using PEI as previously described. 24 hours post transfection, expression was induced with DOX, and samples were fixed in 4% PFA or lysed in 100ul RLT buffer at 0, 30, 60, 120 min post DOX. To test *kaposin* blockade during infection, WT BAC16 or naïve iSLK cells were reactivated with DOX 24 hours after ASO transfection and samples were fixed in 4% PFA or lysed in 100ul RLT buffer at 0, 4, 8 and 24h post DOX for viral transcript analysis. RNA was extracted using the RNeasy Plus Mini Kit (Qiagen) and contaminating DNA was removed using TURBO DNase as per manufacturer’s instructions. qPCR to assess viral gene expression was performed as described below (PCR and RT-qPCR). qPCR reactions were prepared in SsoFast EvaGreen Mastermix as per manufacturer’s instructions, using 2.5 μL of diluted cDNA per reaction and final primer concentrations of 500 nM. qPCR for *MALAT1* utilized specific primers and pCDH-MALAT1 to generate a standard curve of *MALAT1* transcript copies.

For *kaposin* transcript stability (*n* = 3) was analyzed using a two-way repeated measures ANOVA test followed by a Tukey’s post-hoc test (*p* < 0.05). To apply statistical stringency on *n* = 2 experiments testing MALAT1 degradation, data was analyzed using a mixed-effects analysis with a Bonferroni’s multiple comparisons test. This adjusts the *p*-value by the number of comparisons made, with significance at *p*_adj_ < 0.0125 for 293Ts and *p*_adj_ < 0.01 for iSLKs. Data from viral gene expression was analyzed using a two-way repeated measures ANOVA test followed by a Dunnett’s post-hoc test (*p* < 0.05).

### RNA structure prediction

RNA structure predication was performed using the software RNAstructure ^67^(Version 6.5). Briefly, the nucleotide sequences for indicated RNAs were entered into the Single Sequence secondary structure prediction tool. MEA (MaxExpect) structures were predicated, uploaded into the Structure Editor, and basepair probabilities were colour-coded using the associated Partition Save File (*pfs*).

### PCR and RT-qPCR

RNA was collected using either the RNeasy Plus Mini Kit (Qiagen) or TRIzol extraction according to the manufacturer’s instructions and stored at −80°C until further use. RNA concentration was determined using NanoDrop OneC (ThermoFisher) and 500 ng of RNA was reverse transcribed using MaximaH (ThermoFisher) with a combination of random hexamer and oligo dT primers, according to the manufacturer’s instructions. For qPCR, cDNA was diluted 1in10 and SsoFast EvaGreen Mastermix (Biorad) was used for amplification. The ΔΔquantitation cycle (Cq) method was used to determine the fold change in expression of target transcripts using either 18S or HPRT as a housekeeping control gene. For PCR, Taq 2X Mastermix (NEB) and gene specific primers were used to amplify cDNA and products were resolved using a 2% agarose gel and imaged via the ChemiDoc Touch Imaging system (BioRad). Primer sequences for both qPCR and PCR can be found in Table 9 and Table 10, respectively.

**Table 9:**
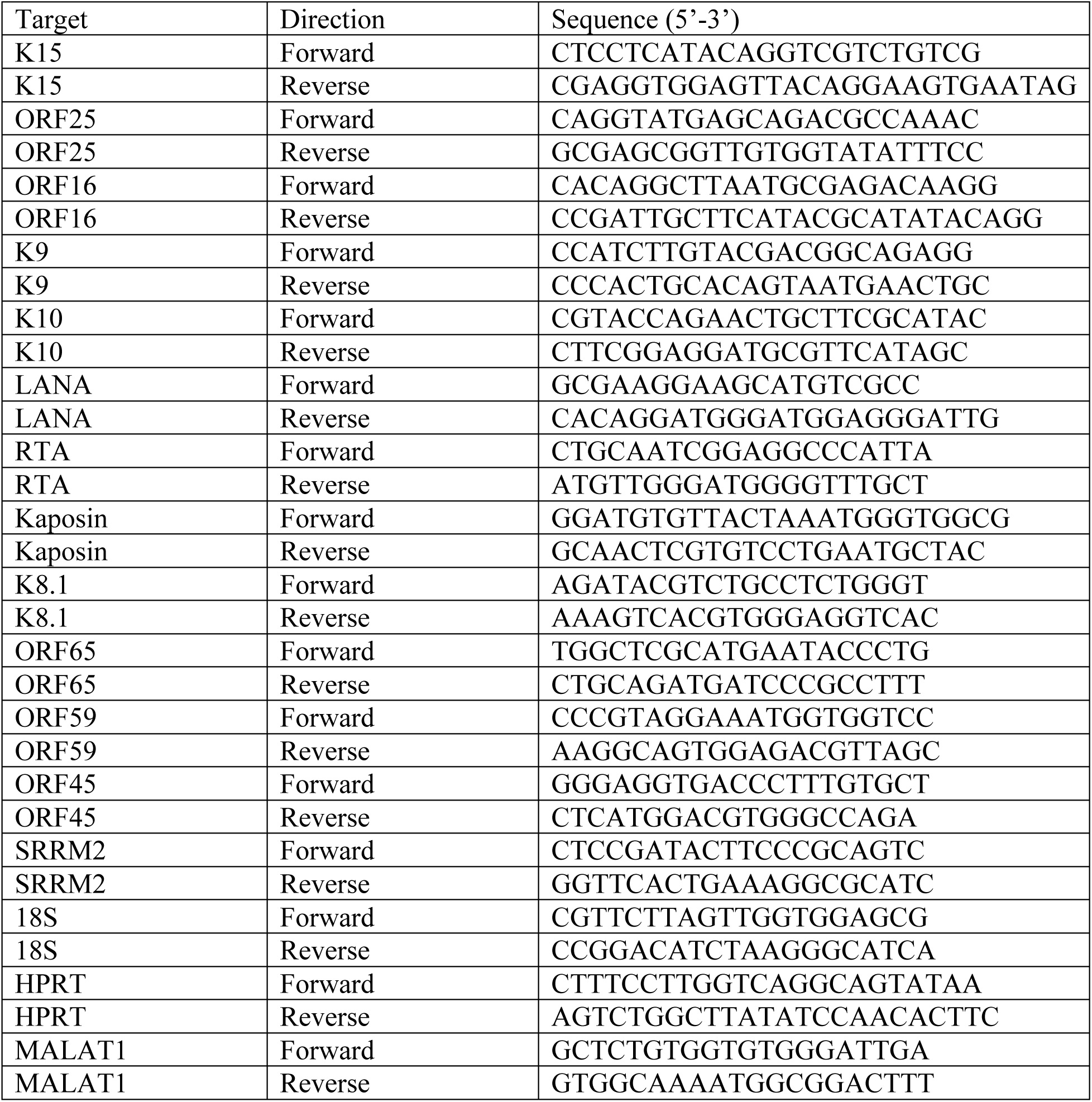
qPCR primers:

**Table 10:**
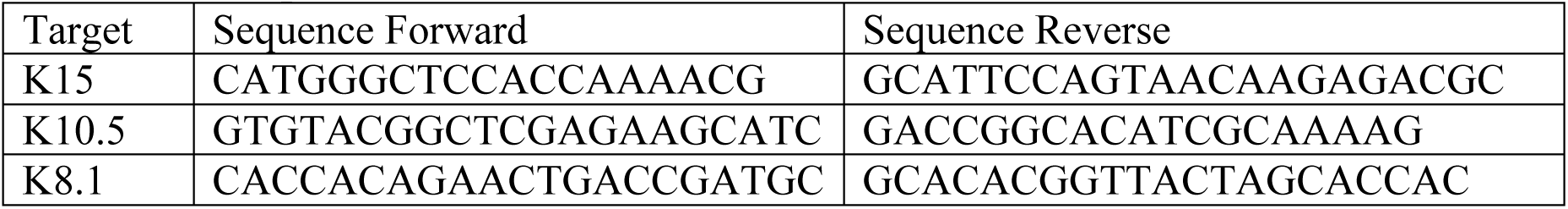
PCR primers.

### TaqMan PCR analysis for transcript copy number

12-well dishes of transfected 293Ts or 48 hour-reactivated WT BAC16/naïve iSLKs were washed in PBS and lysed in 100 ul of RLT buffer for total transcript analysis. In parallel, cell were counted from matched wells. RNA was extracted using the RNeasy Plus Mini Kit as per manufacturer’s instructions, including an on-column DNase I (ThermoFisher, Cat# EN0521) digestion step. Probe-based qPCR using a custom kaposin-K12 TaqMan assay was conducted as above (nuclear/cytoplasmic fractionation), using 500 ng of RNA for cDNA synthesis and cDNA diluted 1:10 for qPCR reactions, meaning each qPCR reaction is estimated represent 12.5 ng of RNA. To convert this into the number of cells represented in the reaction, total RNA yield was first calculated by multiplying RNA concentration with its elution volume. The hemacytometer cell counts were then divided by the matched sample RNA yield to estimate the number of cells represented per unit RNA; when this is multiplied by the amount of RNA represented in the reaction (i.e., 12.5 ng), the product is then the number of cells represented in the reaction.

Resulting qPCR reaction TCN was divided by the number of cells represented in each reaction to approximate the number of *kaposin* transcripts detected on a per cell basis within a sample.

### Lentivirus production and transduction

All recombinant lentiviruses were generated using a second-generation system. 293T cells were transfected with psPAX2, MD2-G, and the lentiviral transfer plasmid pLJM1 or pLKO containing either a gene interest or shRNA (Table 6), respectively, using polyethylenimine (PEI, Polysciences). 4 h post-transfection, serum-free media was replaced with DMEM containing serum but no antibiotics. Viral supernatants were harvested 48 h post-transfection and stored at −80°C prior to use. For transduction, lentiviruses were thawed at 37°C and added to target cells in complete media containing 5 μg/mL polybrene (Sigma-Aldrich). After 24 hours, the media was replaced with selection media containing 5 μg/mL blasticidin (ThermoFisher) and cells were selected for 48 h before proceeding with experiments.

### Immunoprecipitation and Sample Preparation

15-cm dishes of 80-90 percent confluent naïve or BAC16 infected iSLK cells were induced for reactivation with doxycycline. After 48 hours, cells were trypsinized, washed with ice-cold PBS, and resuspended in 500ul of IP-MS Cell Lysis buffer from the Pierce MS-compatible magnetic IP kit (Protein A/G) (Thermo Fisher Scientific Cat#90409). Lysis buffer was supplemented with protease inhibitors (Sigma), sodium fluoride, and sodium orthovanadate. Cell lysates were sonicated using a Bioruptor UCD-200 sonicator 30s ON/90s off, for 1 hour. Cellular debris was removed by centrifugation at 21000 rcf for 10 minutes at 4C. Supernatants were transferred to a fresh tube. 25ul was reserved, and the remaining 475ul of each sample was incubated with either 2.5 ul of SC35 mAb (mouse ascites, Sigma Aldrich, S4045) or 2.5 ul of control mouse IgG1 (Cell Signaling Technologies). Immune complexes were allowed to form overnight in the cold room with constant rotation. The next morning, 25ul pf Protein A/G magnetic beads were washed IP-MS Cell Lysis buffer incubated with lysate for 1-2 hours in the cold room with constant rotation. Beads were then washed with 500 µL of ice-cold 50 mM Tris.Cl pH 7.4, 100 mM NaCl, 0.1% Tween-20 then resuspended with the same buffer supplemented with RNaseI (Ambion, AM2295, final concentration 0.02 U/µL) and incubated at 37°C for 5 minutes according to the protocol in ^11^. The beads were then washed three times with ‘Wash A (+10 mM MgCl2)’ and twice with ‘Wash B’ buffer.

### On Bead Digest and Mass Spectrometry Analysis

The beads were washed with 250 µl 50mM ammonium bicarbonate then resuspended in 100 µl for reduction and alkylation. Cysteines were reduced in 5 mM Tris (2-carboxyethyl) phosphine with shaking, for 20 minutes followed by alkylation in 55 mM chloracetamide for 20 minutes in the dark. Protein was digested on-bead overnight at 37°C, with shaking, using 0.25 µg of MS-grade trypsin (Pierce, Catalog # 90057). The beads were centrifuged (12000xg for 5 minutes) and the supernatant transferred to a new tube. The peptides were dried down in a vacuum centrifuge and resuspended in 0.1% formic acid for desalting using a C18 ZipTip (Millipore Sigma, ZTC185096). The recovered peptides were resuspended in 20 μL of 0.1% formic acid and injected onto a Vanquish Neo UHPLC coupled to an Orbitrap Astral mass spectrometer. The sample was loaded onto 5 cm x 100 μm ID NanoShield C18 trap column (C18, 3μm particle size, 120 Å pore size, Ion Opticks) and eluted onto a 15 cm x 75 μm ID Aurora Elite XT column (C18, 1.7 μm particle size, IonOpticks). The flow rate was 400 nL/min with a column temperature of 40 °C. Mobile phase A consisted of 0.1% (v/v) formic acid in water and mobile phase B consisted of 0.1% (v/v) formic acid in 80:20 acetonitrile: water (v/v). Peptides were separated using a 50-minute linear gradient from 2-40% B.

The Orbitrap Astral mass spectrometer was operated with a resolution of 120,000 (MS^1^) for *m/z* 380–1400. The Orbitrap AGC was set to 300% with a maximum injection time (maxIT) of 3 ms. DIA MS^2^ scans were acquired with the Astral analyzer for *m/z* 380–980 with a normalized AGC target of 500% and a maxIT of 3 ms. Isolation windows were set to 4 Th with 27% Normalized Collision Energy for HCD fragmentation. Raw data files were analyzed in DIA-NN 2.2.0 ^90^ using an *in-silico* DIA-NN predicted spectral library (11,194,365 precursors, allowing for C carbamidomethylation, N-terminal M excision, N-terminal acetylation, M oxidation and 2 missed cleavages). The spectral library was generated from combined human (UP000005640, 20652 sequences) and Human herpesvirus 8 (UP000141755; 87 sequences) reference databases. The search included the following additional settings: Protein inference = ‘Peptidoform’, Neural network classifier = ‘Single-pass mode’, Quantification strategy = ‘Robust LC (high precision)’, Cross-run normalization = ‘RT-dependent’, Library Generation = ‘IDs, RT and IM Profiling’.

Mass accuracy and MS^1^ accuracy were set to 0 for automatic inference and ‘MBR’ was checked. Protein identifications with q-value less than 1% were accepted as determined via the software’s target-decoy strategy.

The mass spectrometry proteomics data have been deposited to the ProteomeXchange Consortium via the PRIDE partner repository with the dataset identifier PXD080367. **RNA-seq** RNA from 5 replicates of BAC16-iSLK cells, either latent or 48 hrs post-reactivation, was collected using the RNeasy Plus Mini Kit (Qiagen) according to standard protocol. RNA was quantified using the Qubit™ RNA Broad Sensitivity Assay Kit (Thermo Fisher) and a Qubit 4 Fluorometer (Invitrogen). At least 10 ng of RNA/sample was submitted for library preparation and sequencing at the University of Calgary Center for Health Genomics and Informatics.

Libraries were prepared using the NEBNext Ultra II RNA kit (NEB) with rRNA depletion (NEB) and samples were sequenced on one NovaSeq S1 200 cycle v1.5 run with 200 cycles (bp) for a total output of ∼32M read pairs per sample. GEO accession numbers for the data set can be found in Table 12.

**Table 11:**
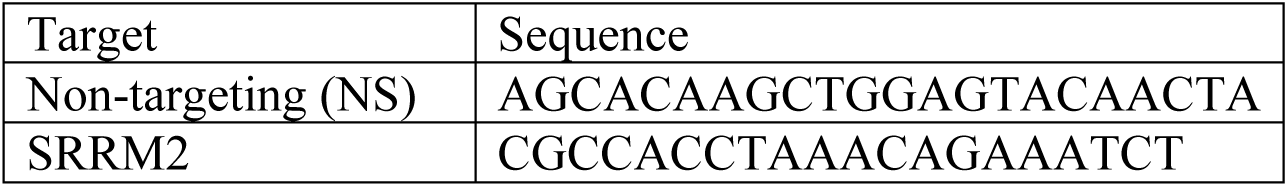
shRNA sequences.

**Table 12:** GEO Accession Numbers for RNA sequencing.

| Accession Number | Sample Name | Biological Replicate |
| --- | --- | --- |
| GSM8543561 | Mini DRs Lytic 48h | 1 |
| GSM8543562 | Mini DRs Lytic 48h | 2 |
| GSM8543563 | Mini DRs Lytic 48h | 3 |
| GSM8543564 | Mini DRs Lytic 48h | 4 |
| GSM8543565 | Mini DRs Lytic 48h | 5 |
| GSM8543566 | Mini DRs Latent | 1 |
| GSM8543567 | Mini DRs Latent | 2 |
| GSM8543568 | Mini DRs Latent | 3 |
| GSM8543569 | Mini DRs Latent | 4 |
| GSM8543570 | Mini DRs Latent | 5 |
| GSM8543571 | ΔB Lytic 48h | 1 |
| GSM8543572 | ΔB Lytic 48h | 2 |
| GSM8543573 | ΔB Lytic 48h | 3 |
| GSM8543574 | ΔB Lytic 48h | 4 |
| GSM8543575 | ΔB Lytic 48h | 5 |
| GSM8543576 | ΔBC Lytic 48h | 1 |
| GSM8543577 | ΔBC Lytic 48h | 2 |
| GSM8543578 | ΔBC Lytic 48h | 3 |
| GSM8543579 | ΔBC Lytic 48h | 4 |
| GSM8543580 | ΔBC Lytic 48h | 5 |
| GSM8543581 | ΔBC Latent | 1 |
| GSM8543582 | ΔBC Latent | 2 |
| GSM8543583 | ΔBC Latent | 3 |
| GSM8543584 | ΔBC Latent | 4 |
| GSM8543585 | ΔBC Latent | 5 |
| GSM8543586 | ΔB Latent | 1 |
| GSM8543587 | ΔB Latent | 2 |
| GSM8543588 | ΔB Latent | 3 |
| GSM8543589 | ΔB Latent | 4 |
| GSM8543590 | ΔB Latent | 5 |
| GSM8543591 | ΔC Lytic 48h | 1 |
| GSM8543592 | ΔC Lytic 48h | 2 |
| GSM8543593 | ΔC Lytic 48h | 3 |
| GSM8543594 | ΔC Lytic 48h | 4 |
| GSM8543595 | ΔC Lytic 48h | 5 |
| GSM8543596 | ΔC Latent | 1 |
| GSM8543597 | ΔC Latent | 2 |
| GSM8543598 | ΔC Latent | 3 |
| GSM8543599 | ΔC Latent | 4 |
| GSM8543600 | ΔC Latent | 5 |
| GSM8543601 | WT Lytic 48h | 1 |
| GSM8543602 | WT Lytic 48h | 2 |
| GSM8543603 | WT Lytic 48h | 3 |
| GSM8543604 | WT Lytic 48h | 4 |
| GSM8543605 | WT Lytic 48h | 5 |
| GSM8543606 | WT Latent | 1 |
| GSM8543607 | WT Latent | 2 |
| GSM8543608 | WT Latent | 3 |
| GSM8543609 | WT Latent | 4 |
| GSM8543610 | WT Latent | 5 |

**Table 13:**
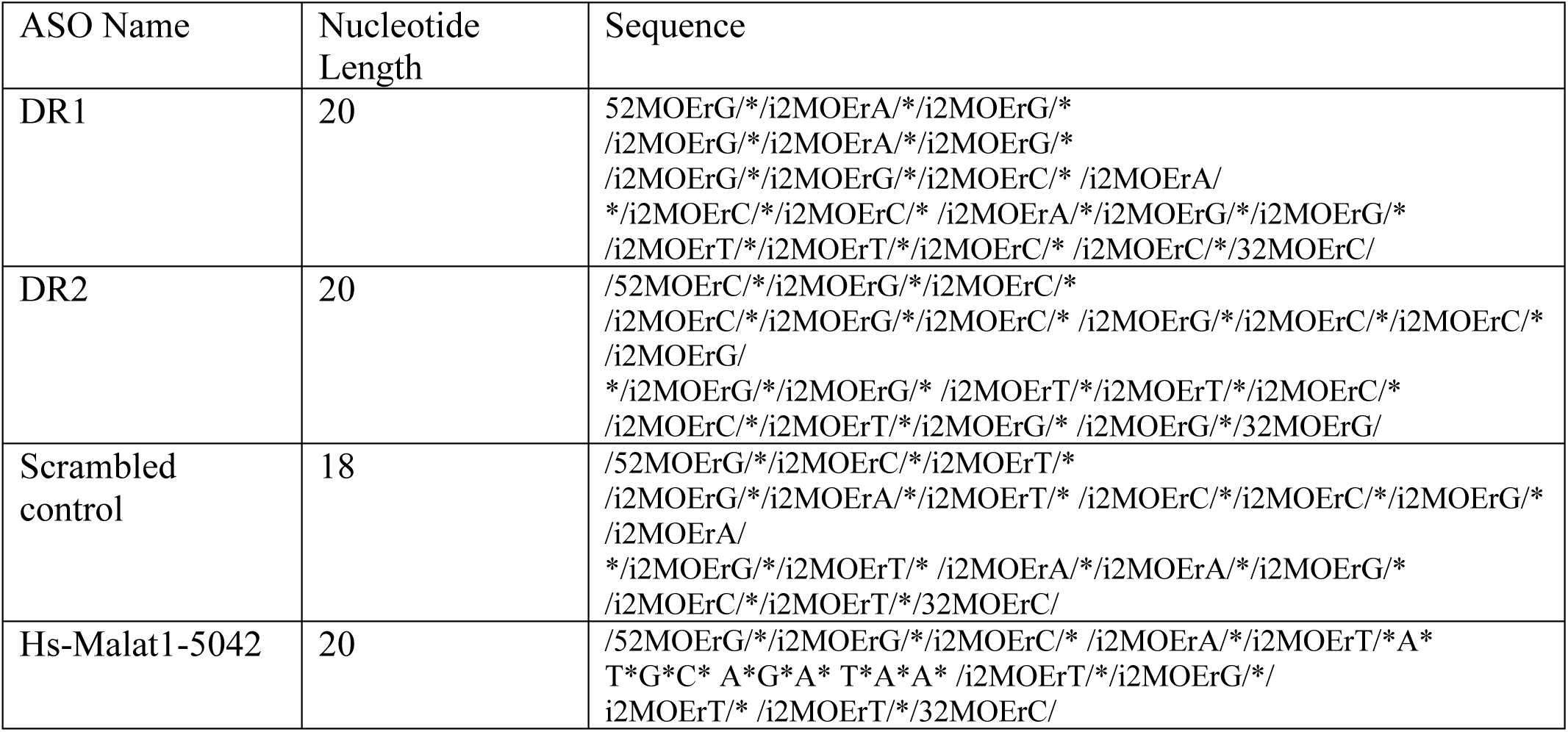
Antisense Oligonucleotides.

### RNA-seq analysis

#### Quality control

Quality of the RNA-Seq libraries and sequencing runs was verified using FastQC (v0.12.1)^91^. FastQC: a quality control tool for high throughput sequence data. Available online at: http://www.bioinformatics.babraham.ac.uk/projects/fastqc]).

#### Reference genome

A custom reference genome was generated to include both human and KSHV sequences. The FASTA sequences for Homo sapiens reference GCA_000001405.15_GRCh38_full_plus_hs38d1_analysis_set and Human herpesvirus 8 strain JSC-1 clone BAC16 (GenBank GQ994935.1 were concatenated to generate “human_plus_kshv.fa”.

Reference gene annotations were combined from GENCODE human v43, GenBank (KSHV; GQ994935.1), and three additional entries corresponding to kaposin mRNA, PAN lncRNA, and T1.4 RNA to generate “human_plus_kshv.gff3”.

#### Differential expression testing

The Salmon (v1.10.2)^92^ function ‘index’ was used to generate the salmon index, with genomic sequences considered to be decoys for the transcriptome alignments. RNA-Sequencing reads were subsequently quantified using ‘salmon quant’ pseudoalignment with parameters ‘--libType ISR’. Count tables were generated using tximport (v1.28.0)^93^. Transcript-level counts used parameters ‘type = salmon, dropInfReps = TRUE, txOut = TRUE, countsFromAbundance = dtuScaledTPM’. Gene-level counts used parameters ‘type = salmon, dropInfReps = TRUE, txOut = FALSE, countsFromAbundance = lengthScaledTPM’. Differential expression testing was performed on both transcripts and genes using limma (v3.54)^94^ with voom transformation and Benjamini-Hochberg correction applied to the p-values.

Heat map was generated using log2FC values.

### Image analysis

Nuclear speckles were quantified using an unbiased image analysis pipeline generated in the freeware CellProfiler4.0.6 (cellprofiler.org) in a method similar to ^50,53,95^. First, nuclear staining was used to identify individual cells by applying a threshold to remove background staining and the “IdentifyPrimaryObjects” module for objects between 150 and 400 pixels in diameter. For each identified object (nuclei), the peripheral boundary (cytoplasm) of each cell was defined using the “IdentifySecondaryObjects” module by expanding a distance of 150 pixels from the center of each nucleus and the subsequent objects defined in this step were named “cells”.

Importantly, multiple nuclei in proximity (e.g., within the propagation distance) would be divided into mutually exclusive cells, such that a single NS could not be counted more than once. NSs in the entire field of view were identified by applying a threshold to remove background staining and the “IdentifyPrimaryObjects” module for objects of between 6 and 50 pixels in diameter. The output of this module was named “NS in all cells”. To quantify NSs in *kaposin*-positive cells only, *kaposin* staining was first identified by applying a threshold to remove background staining and then used as an inverse mask for objects previously defined as “cells”. In doing so, any cell that did not stain positive for *kaposin* was removed or “masked” from the remainder of the analysis. NSs were then identified as before, and the output of this module was named “NS in *kaposin*-positive cells only”. The intensity, size, and eccentricity of the identified NSs were determined using the “MeasureObjectIntensity” and “MeasureObjectSizeShape” modules and values from the “AreaShape_Area”, “Intensity_IntergratedIntensity” and “Area_Eccentricity” categories were graphed for analysis. To measure colocalization of *kaposin*-puncta, NSs, and/or *MALAT1* either *kaposin*-puncta or NSs were first identified by applying a threshold to remove background staining and the “IdentifyPrimaryObjects” module for objects of between 8 and 50 pixels in diameter. The “MeasureObjectColocalization” module was used to measure colocalization between the identified objects and the stain of interest (for example, if NSs were identified then the colocalization of staining from either the *kaposin* or *MALAT1* channel was quantified) and values from the “Correlation_Correlation” category were graphed for analysis. For ASO experiments in Figures 5 and Suppl Fig 7, images were acquired by 40x-oil widefield microscopy, and the pipeline was modified. Nuclei size ranged between 60 to 220 pixels and *kaposin* NSs size ranged between 3 to 40 pixels.

### Statistics

Statistics were performed using Graphpad Prism version 9 or 10.2.2 except for those specific to RNA-seq which were performed separately. The specific statistical test is specified in the figure legend.

### Graphing

All graphing was done using Graphpad Prism version 9 or 10.2.2.

| REAGENT or RESOURCE | SOURCE | IDENTIFIER |
| --- | --- | --- |
| **Antibodies** |  |  |
| Mouse monoclonal anti-SRRM2 (referred to as “SC35 mAb” in IP protocol) | Sigma-Aldrich | Cat#S4045 |
| Mouse IgG1 isotype control | Cell Signaling Technology | Cat#5414 |
| Mouse monoclonal anti-SRRM2 | Abcam | Cat#ab11826 |
| Rabbit polyclonal anti-SON | Abcam | Cat#ab121759 |
| Mouse monoclonal anti-PML | Santa Cruz Biotechnology | Cat#sc-966 |
| Mouse monoclonal anti-Coilin | Abcam | Cat#ab87913 |
| Rabbit polyclonal anti-PRPF8 | Abcam | Cat#ab79237 |
| Rabbit polyclonal anti-LANA | Gift from Dr. Craig McCormick (Dalhousie University) | N/A |
| Mouse monoclonal anti-ORF57 | Santa Cruz Biotechnology | Cat#sc-135746 |
| Rabbit polyclonal anti-ORF6 | Gift from Dr. John Karijovich | N/A |
| Rabbit monoclonal anti-Tubulin | Cell Signaling Technology | Cat#2128S |
| Rabbit monoclonal anti-Lamin A/C | Cell Signaling Technology | Cat#2032S |
| Rabbit polyclonal anti-Kaposin B/C | Gift from Dr. Craig McCormick (Dalhousie University) | N/A |
| Chicken anti-Rabbit Alexa Fluor™ 488 (secondary) | Thermo Fisher Scientific | Cat#A21441 |
| Goat anti-Mouse Alexa Fluor™ 488 (secondary) | Thermo Fisher Scientific | Cat#A11029 |
| Donkey anti-Rabbit Alexa Fluor™ 405 (secondary) | Abcam | Cat#ab175651 |
| **Bacterial and virus strains** |  |  |
| One Shot™ Stbl3™ Chemically Competent E. coli | Thermo Fisher Scientific | Cat#C737303 |
| KSHV: BAC16 (wild type) | Brulois et al. | GenBank: GQ994 935.1 |
| KSHV: BAC16-ΔKapB | Kleer et al. | N/A |
| KSHV: BAC16-ΔKapC | Kleer et al. | N/A |
| KSHV: BAC16-ΔKapBC | Kleer et al. | N/A |
| KSHV: BAC16-miniDRs | This paper | N/A |
| **Experimental models: Cell lines** |  |  |
| HEK293T | ATCC | Cat#CRL-3216 |
| HeLa | ATCC | N/A |
| Naïve iSLK-TREx-RTA | Myoung and Ganem | N/A |
| iSLK-TREx-RTA stably infected with KSHV BAC16 (WT) | Kleer et al. | N/A |
| iSLK-TREx-RTA stably infected with KSHV BAC16-ΔKapB (ΔKapB iSLK) | Kleer et al. | N/A |
| iSLK-TREx-RTA stably infected with KSHV BAC16-ΔKapC (ΔKapC iSLK) | Kleer et al. | N/A |
| iSLK-TREx-RTA stably infected with KSHV BAC16-ΔKapBC (ΔKapBC iSLK) | Kleer et al. | N/A |
| iSLK-TREx-RTA stably infected with KSHV BAC16-ΔKapABC (ΔKapABC iSLK) | Kleer et al. | N/A |
| iSLK-TREx-RTA stably infected with KSHV BAC16-miniDRs | This paper | N/A |
| BCBL1-TREx-RTA | Gift from Dr. Craig McCormick | N/A |
| HUVEC | Lonza | Cat#C2519A |
| **Chemicals, peptides, and recombinant proteins** |  |  |
| Poly-D-lysine | Sigma-Aldrich | Cat#A3890401 |
| Avidin, NeutrAvidin™ Oregon Green™ 488 conjugate | Thermo Fisher Scientific | Cat#A6374 |
| Chloramphenicol | Sigma-Aldrich | Cat#C0378-5G |
| Geneticin | Thermo Fisher Scientific | Cat# G8168-10ML |
| Hygromycin B | Thermo Fisher Scientific | Cat#10687010 |
| SUPERase•In™ RNase Inhibitor | Thermo Fisher Scientific | Cat#AM2694 |
| RNaseI | Ambion | Cat#AM2295 |
| Trypsin (MS-grade) | Pierce (Thermo Fisher Scientific) | Cat#90057 |
| Actinomycin D | Sigma-Aldrich | Cat#A9415-2MG |
| Fibronectin | Sigma-Aldrich | Cat#CLS356008 |
| **Critical commercial assays** |  |  |
| RNeasy Plus Mini Kit | Qiagen | Cat#74134 |
| Large-Construct Kit | Qiagen | Cat#12462 |
| TURBO DNA-free™ Kit (TURBO DNase) | Thermo Fisher Scientific | Cat#AM2238 |
| DC Protein Assay Kit | Bio-Rad | Cat#5000112 |
| Pierce MS-Compatible Magnetic IP Kit (Protein A/G) | Thermo Fisher Scientific | Cat#90409 |
| Qubit™ RNA Broad Range Assay Kit | Thermo Fisher Scientific | Cat#Q10210 |
| NEBNext Ultra II RNA Library Prep Kit | New England Biolabs | Cat#E7770S |
| NEBNext rRNA Depletion Kit | New England Biolabs | Cat#E7400S |
| BioNick™ DNA Labeling System | Thermo Fisher Scientific | Cat#18247015 |
| Click-iT™ RNA Alexa Fluor™ 488 Imaging Kit | Invitrogen | Cat#C10329 |
| Nuclei EZ Prep Nuclei Isolation Kit | Sigma-Aldrich | Cat#NUC-101 |
| **Deposited data** |  |  |
| RNA-seq data: BAC16-iSLK WT, ΔKapB, ΔKapC, ΔKapBC and miniDRs, latent and 48 h reactivated | This paper | GSM8543561-GSM8543597 |
| SRRM2 immunoprecipitation – mass spectrometry (IP-MS) proteomics data from naïve and reactivated iSLK-BAC16 WT and ΔKapB cells | This paper | ProteomeXchange, identifier PXD080367 |
| **Recombinant DNA** |  |  |
| pcDNA3.1(+) | Invitrogen | Cat#V79020 |
| pcDNA-kaposin | BioBasic (custom gene synthesis) | N/A |
| pcDNA-kaposin-reverse | This paper (cloned into pcDNA3.1 using NheI and BamHI) | N/A |
| pcDNA-kaposin-ΔStarts | BioBasic (custom gene synthesis) | N/A |
| pcDNA-kaposin-altKapB [N7] | BioBasic (custom gene synthesis) | N/A |
| pcDNA-kaposin-ΔStarts-ΔDR1 | BioBasic (custom gene synthesis) | N/A |
| pcDNA-kaposin-ΔStarts-ΔDR2 | BioBasic (custom gene synthesis) | N/A |
| pcDNA-kaposin-ΔStarts-DRswap | BioBasic (custom gene synthesis) | N/A |
| pcDNA-kaposin-miniDR | BioBasic (custom gene synthesis) | N/A |
| pcDNA-kaposin-miniDR-swap | BioBasic (custom gene synthesis) | N/A |
| pcDNA-kaposin-5X-DR1 | BioBasic (custom gene synthesis) | N/A |
| pcDNA-kaposin-3X-DR2 | BioBasic (custom gene synthesis) | N/A |
| pcDNA-rtTA | This paper (cloned into pcDNA3.1 using NheI and NotI) | N/A |
| MS2_GFP | Gift from Daniel Larson | Addgene #61764 |
| MCP-NLS-EGFP | This paper (cloned from MS2_GFP using BamHI and NotI) | N/A |
| pEGFP-SF2 | Gift from Tom Misteli | Addgene #17990 |
| mCherry-SRSF1 | This paper (cloned from pEGFP-SF2 using HindIII and BamHI) | N/A |
| mCherry-RBM25 | This paper (cloned from cDNA using NheI and SalI) | N/A |
| pTRE-SOX | Gift from Dr. Eric Pringle | N/A |
| pcDNA-puro-TagBFP-β-ACTIN-3'UTR-24xMS2 | Gift from Weirui Ma | Addgene #205161 |
| pTRE-EV-MS2 | This paper (cloned from pTRE-SOX and pcDNA-puro-TagBFP-β-ACTIN-3'UTR-24xMS2) | N/A |
| pTRE-kaposin-ΔStarts-MS2 | This paper (cloned into pTRE-EV-MS2 from pcDNA-kaposin-ΔStarts using NheI and EcoRI) | N/A |
| pTRE-kaposin-ΔDR1-MS2 | This paper (cloned into pTRE-EV-MS2 from pcDNA-kaposin-ΔStarts-ΔDR1 using NheI and EcoRI) | N/A |
| pTRE-kaposin-ΔDR2-MS2 | This paper (cloned into pTRE-EV-MS2 from pcDNA-kaposin-ΔStarts-ΔDR2 using NheI and EcoRI) | N/A |
| pTRE-4X-DR2-MS2 | This paper (cloned into pTRE-EV-MS2 using annealed oligos with NheI/EcoRI sticky ends) | N/A |
| pTRE-4X-DR1-MS2 | This paper (cloned into pTRE-EV-MS2 using annealed oligos with NheI/EcoRI sticky ends) | N/A |
| pTRE-4X-DR2-reverse-MS2 | This paper (cloned into pTRE-EV-MS2 using annealed oligos with NheI/EcoRI sticky ends) | N/A |
| pTRE-4X-DR2-scramble-MS2 | This paper (cloned into pTRE-EV-MS2 using annealed oligos with NheI/EcoRI sticky ends) | N/A |
| pTRE-RRV-3X-rDL-E2-MS2 | This paper (cloned into pTRE-EV-MS2 using annealed oligos with NheI/EcoRI sticky ends) | N/A |
| pTRE-RRV-5X-rDL-E1-MS2 | This paper (cloned into pTRE-EV-MS2 using annealed oligos with NheI/EcoRI sticky ends) | N/A |
| pcDNA-ORF57 | BioBasic (custom gene synthesis) | N/A |
| psPAX2 | Gift from Didier Trono | Addgene #12260 |
| pMD2.G | Gift from Didier Trono | Addgene #12259 |
| pLKO.1-blast | Gift from Keith Mostov | Addgene #26655 |
| pLKO-NT (non-targeting shRNA) | Gift from Dr. Craig McCormick (Dalhousie University) | N/A |
| pLKO-SRRM2-sh1 | This paper (cloned into pLKO.1-blast using AgeI and EcoRI) | N/A |
| pLKO-SRRM2-sh2 | This paper (cloned into pLKO.1-blast using AgeI and EcoRI) | N/A |
| pCDH-hMALAT1 | Gift from Jianjun Zhao | Addgene #118580 |
| **Oligonucleotides** |  |  |
| Custom TaqMan® probe/primers: kaposin (KapA region; Assay ID AP7D4GA) | This paper | Assay ID: AP7D4GA |
| qPCR primers (17 targets; see Table 4) | This paper | N/A |
| PCR primers (3 targets; see Table 5) | This paper | N/A |
| Stellaris® FISH Probes, Human MALAT1 with Quasar® 670 Dye | Biosearch Technologies | Cat# VSMF-2211-5 |
| Stellaris® FISH Probes, Human NEAT1_m with Quasar® 670 Dye | Biosearch Technologies | Cat#VSMF-2251-5 |
| Custom Stellaris® FISH probes: kaposin and PAN (see Table 2) | Biosearch Technologies | N/A |
| shRNA target sequences: non-targeting control and SRRM2 (see Table 6) | This paper | N/A |
| Antisense oligonucleotides: kaposin DR1, DR2, scrambled control and MALAT1 control gapmer (see Table 8) | Integrated DNA Technologies (IDT) | N/A |
| **Software and algorithms** |  |  |
| GraphPad Prism (v9 and v10.2.2) | GraphPad Software | https://www.graphpad.com |
| ImageJ | National Institutes of Health | https://imagej.nih.gov/ij/ |
| CellProfiler (v4.0.6) | Broad Institute | https://cellprofiler.org |
| RNAstructure (v6.5) | Reuter and Mathews<sup>67</sup> | https://rna.urmc.rochester.edu/RNAstructure.html |
| DIA-NN (v2.2.0) | Demichev et al<sup>90</sup> | https://github.com/vdemichev/DiaNN |
| FastQC (v0.12.1) | Babraham Bioinformatics | http://www.bioinformatics.babraham.ac.uk/projects/fastqc/ |
| The Salmon (v1.10.2) | Patro et al<sup>92</sup> | https://combine-lab.github.io/salmon/ |
| tximport (v1.28.0) | Soneson et al<sup>93</sup> | https://bioconductor.org/packages/release/bioc/html/tximport.html |
| limma (v3.54) | Ritchie et al<sup>94</sup> | https://bioconductor.org/packages/release/bioc/html/limma.html |

## Supplemental Figures

**Supplemental Figure 1.**
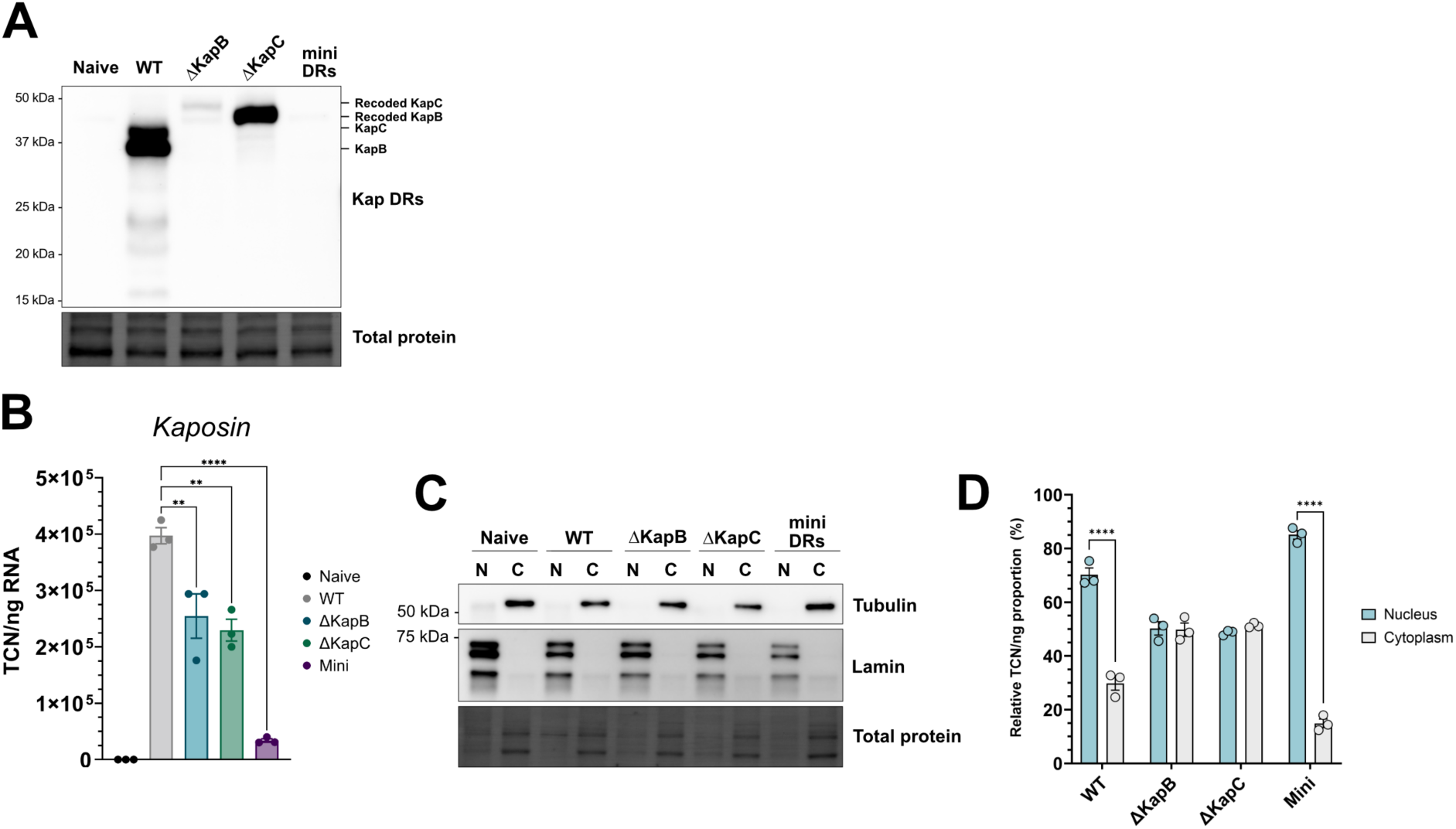
Direct repeat regions retain *kaposin* in the nuclear fraction during infection. (**A-D**) Naïve iSLK or iSLK-BAC16 cells were reactivated for 48h and either lysed for total protein (**A**) or RNA (**B**) or fractionated into nuclear and cytoplasmic fractions before lysis for protein (**C**) or RNA (**D**). (**A, C**) Immunoblots for Kap polypeptides using anti-KapDRs polyclonal antibody (**A**) or nuclear (anti-lamin) and cytoplasmic (anti-tubulin) fractions are shown. KapB and KapC molecular weights are lower after infection with WT virus versus ΔKapB/ΔKapC due to a deletion in the repeat region in WT virus, described in ^53^. (**B, D**) *Kaposin* transcript copy number (TCN) per cell (**B**) or per fraction (expressed as TCN/ng RNA) (**D**) were determined by TaqMan PCR and statistics were performed using an ordinary one-way ANOVA with Tukey’s post-hoc (N = 3, p < 0.05) or using a two-way ANOVA with Tukey’s post-hoc (N = 3, p< 0.05), respectively.

**Supplemental Figure 2.**
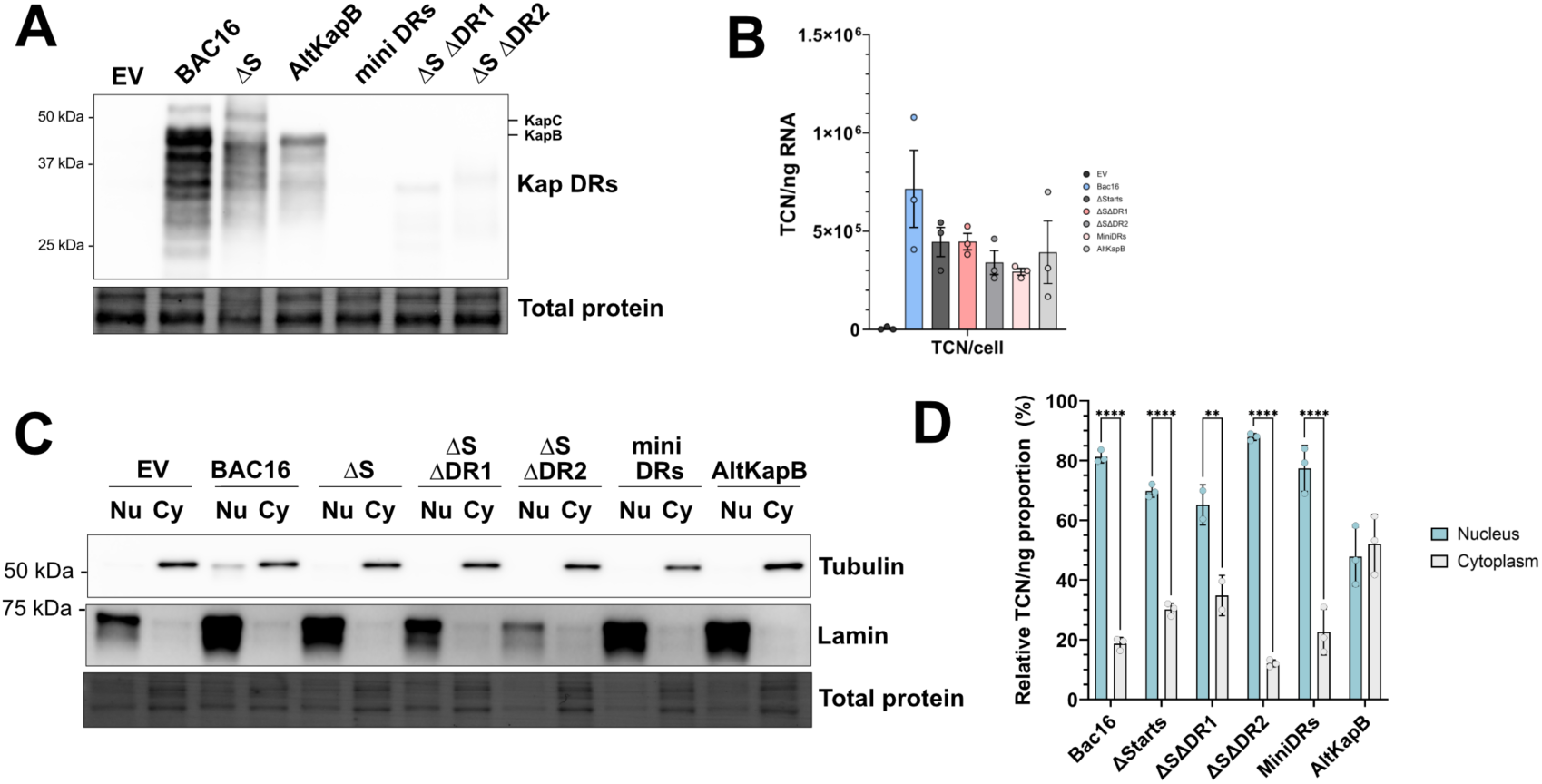
Direct repeat regions retain *kaposin* in the nuclear fraction after transfection. (**A-D**) 293T cells were transfected with the indicated construct for 24 h and either lysed for total protein (**A**) or RNA (**B**) or fractionated into nuclear and cytoplasmic fractions before lysis for protein (**C**) or RNA (**D**). (**A, C**) Immunoblots for Kap polypeptides using anti-KapDRs polyclonal antibody (**A**) or nuclear (anti-lamin) and cytoplasmic (anti-tubulin) fractions are shown. Unexpected Kap polypeptides are translated after transfection of *kaposinΔstarts*, despite lacking all reported start codons. (**B, D**) *Kaposin* transcript copy number (TCN) per cell (**B**) or per fraction (expressed as TCN/ng RNA) (**D**) were determined by TaqMan PCR and statistics were performed using an ordinary one-way ANOVA with Tukey’s post-hoc (N = 3, p < 0.05) or using a two-way ANOVA with Tukey’s post-hoc (N = 3, p< 0.05), respectively.

**Supplemental Figure 3.**
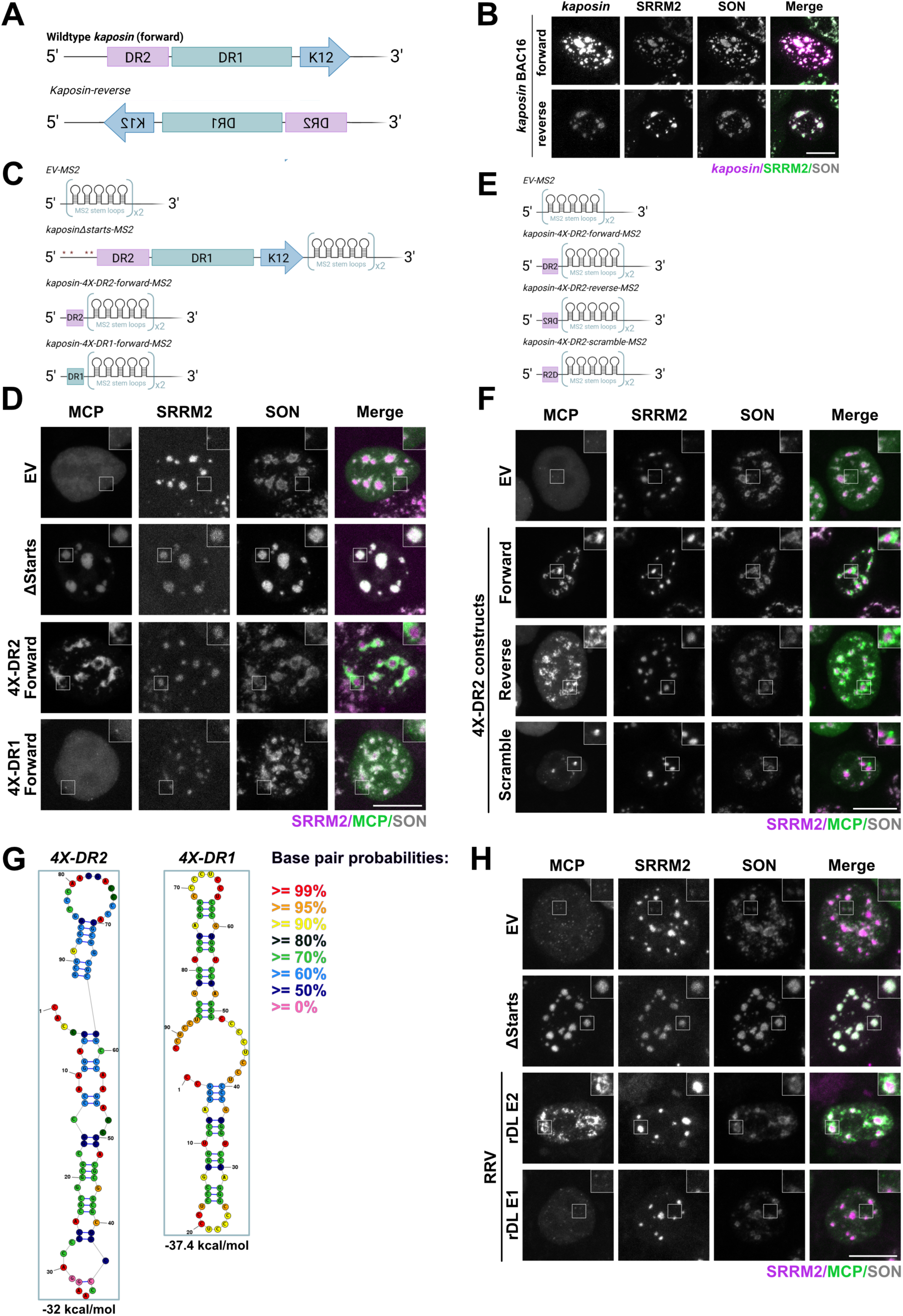
K*a*posin does not require a linear sequence motif for NS co-localization. **(A, C, E)** Diagrams of *kaposin* constructs. A red asterisk indicates a mutated start codon. See Table 5. **(B)** 293T cells were transfected and stained for *kaposin* (magenta), SRRM2 (green), and SON (grey). **(D, F)** 293T cells were transfected with indicated constructs, EGFP-tagged MCP, and rtTA. After induction with Dox, cells were stained for SRRM2 (magenta) and SON (grey). Colour threshold altered for MCP channel for visualization purposes only. **(G)** The secondary structure of *4X-DR2* and *4X-DR1* was modelled using RNAStructure ^81^. Colours correspond to base pairing probabilities. **(H)** 293T cells were transfected with MS2-tagged *EV*, *kaposinΔstarts*, RRV *rDL E2* (3X-E2 repeats), and RRV *rDL E1* (5X-E1 repeats) constructs, EGFP-tagged MCP, and rtTA. After induction with Dox, cells were stained for SRRM2 (magenta) and SON (grey).

**Supplemental Figure 4.**
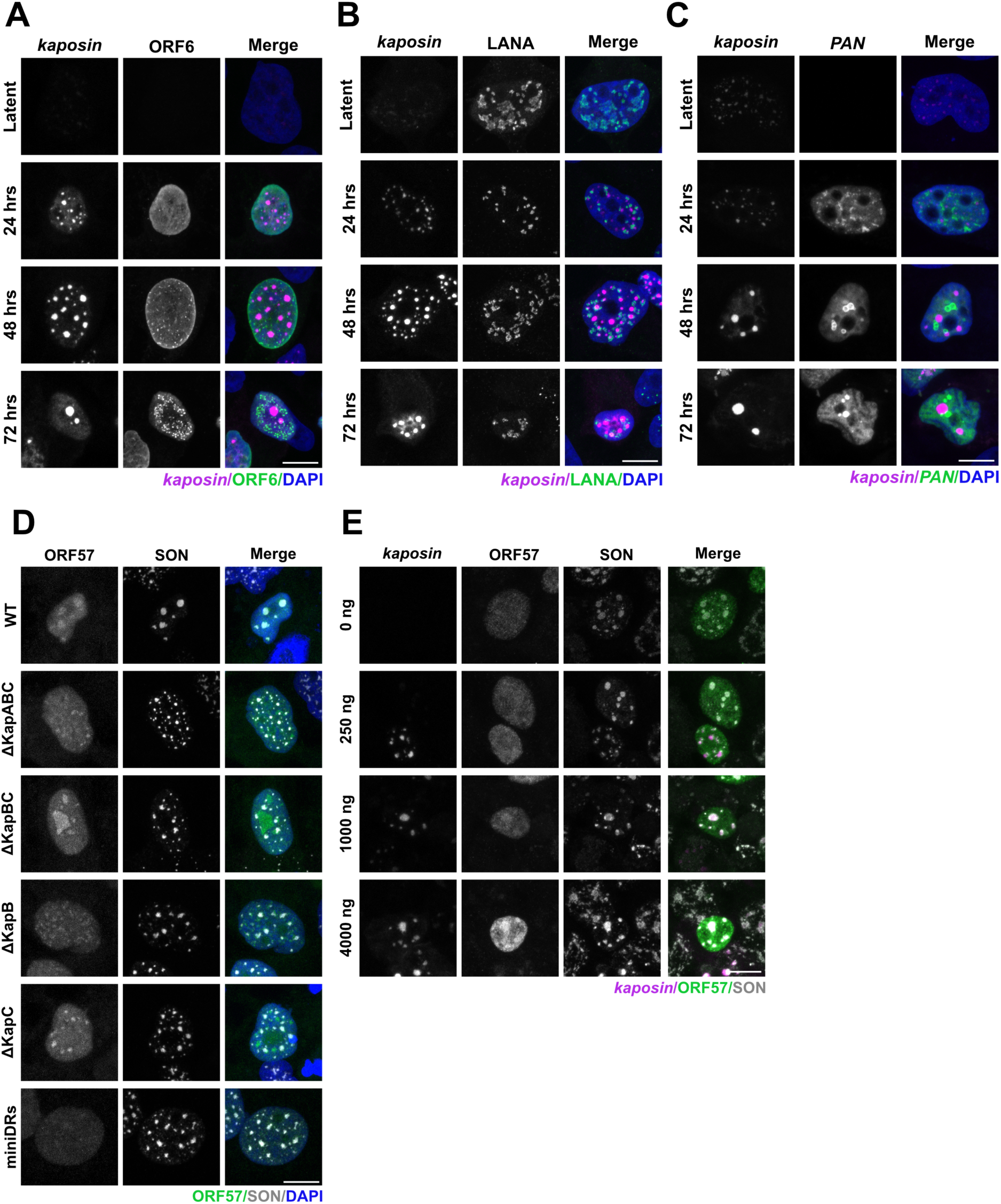
K*a*posin enhances ORF57 recruitment to NSs. **(A-C)** WT iSLK-BAC16s were seeded onto coverslips and treated with Dox for 48 hrs to induce lytic reactivation. Cells were fixed and stained for kaposin (magenta) and either viral ORF6 **(A)**, LANA **(B)**, or the abundant viral lncRNA, PAN **(C)** (green), and nuclei (blue). **(D)** The indicated reactivated iSLK cells were stained for ORF57 (false coloured green) SON (grey), and nuclei (blue). (**E**) 293T cells were transfected with increasing amounts of *kaposinΔstarts* and ORF57 constructs and stained for *kaposin* (magenta), ORF57 (green) and SON (grey).

**Supplemental Figure 5.**
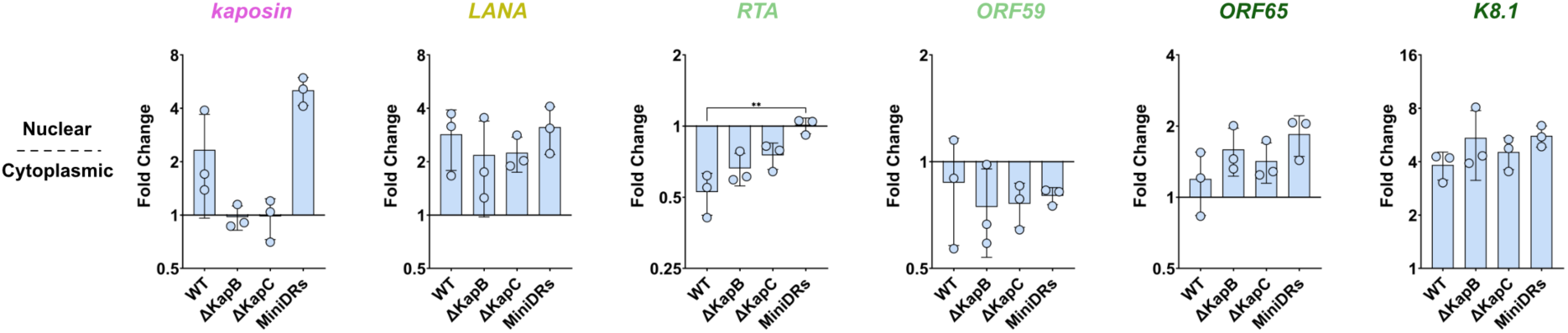
K*a*posin-NS colocalization does not influence viral mRNA export. The indicated reactivated BAC16 iSLK cells were lysed and nuclear and cytoplasmic fractions isolated. RT-qPCR using *kaposin, LANA, RTA*, *ORF59*, *ORF65, K8.1* and *18S* (housekeeping) specific primers was performed. Temporal viral gene class: yellow = latent, light green = early lytic, dark green = late lytic and grey = unclassed, kaposin=pink. Data is presented as fold-change of the cytoplasmic fraction over the nuclear fraction. Statistics were performed using a repeated measures one-way ANVOA with Tukey’s post-hoc (N = 3; *p* < 0.05). Graphs are logscale.

**Supplemental Figure 6.**
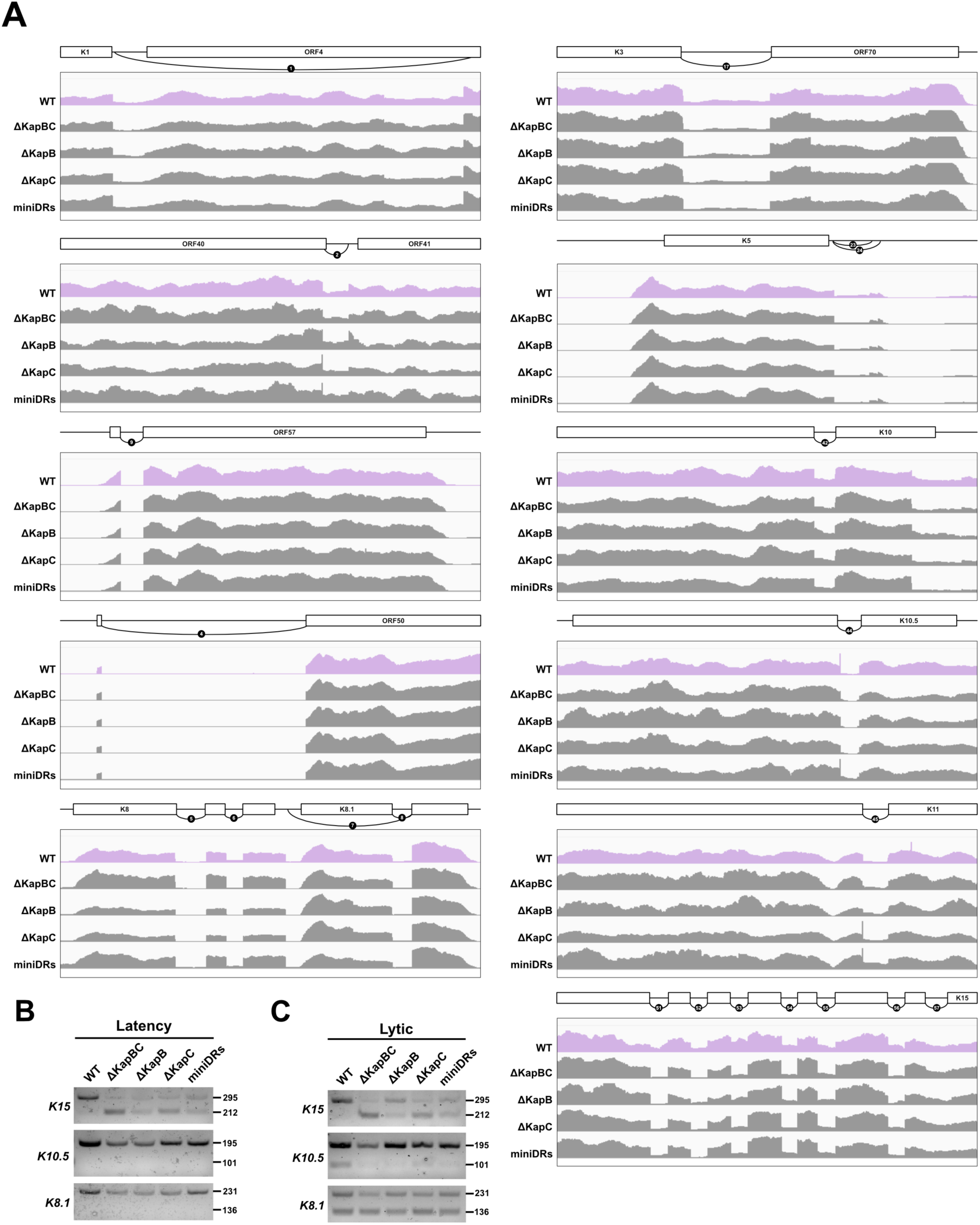
RNA-seq coverage for viral splice junctions. 48h reactivated WT (pink), ΔKapBC, ΔKapB, ΔKapC and miniDRs (grey) iSLK-BAC16 cells showing RNA-seq read coverage of one representative sample (of N=5) for spliced KSHV transcripts. Reads were autoscaled to compensate for differences in transcription and annotated according to (accession #GQ994935). ORFs are indicated with white boxes and previously identified splice junctions are indicted with curves and labelled in accordance with the field. Each “window” spans 3360 bps. **(B-C)** RT-PCR was performed on RNA isolated from the same experiment to examine viral alternative splicing events that may occur during latent **(B)** and lytic **(C)** infection using primers that span splice junctions (SJ) for the following viral genes: *K15* (SJ 53), *K10.5* (SJ 44) and *K8.1* (SJ 8) and cellular genes *ATP6V1D* and *BSCL2*. PCR amplification products were resolved via agarose gel electrophoresis.

**Supplemental Figure 7.**
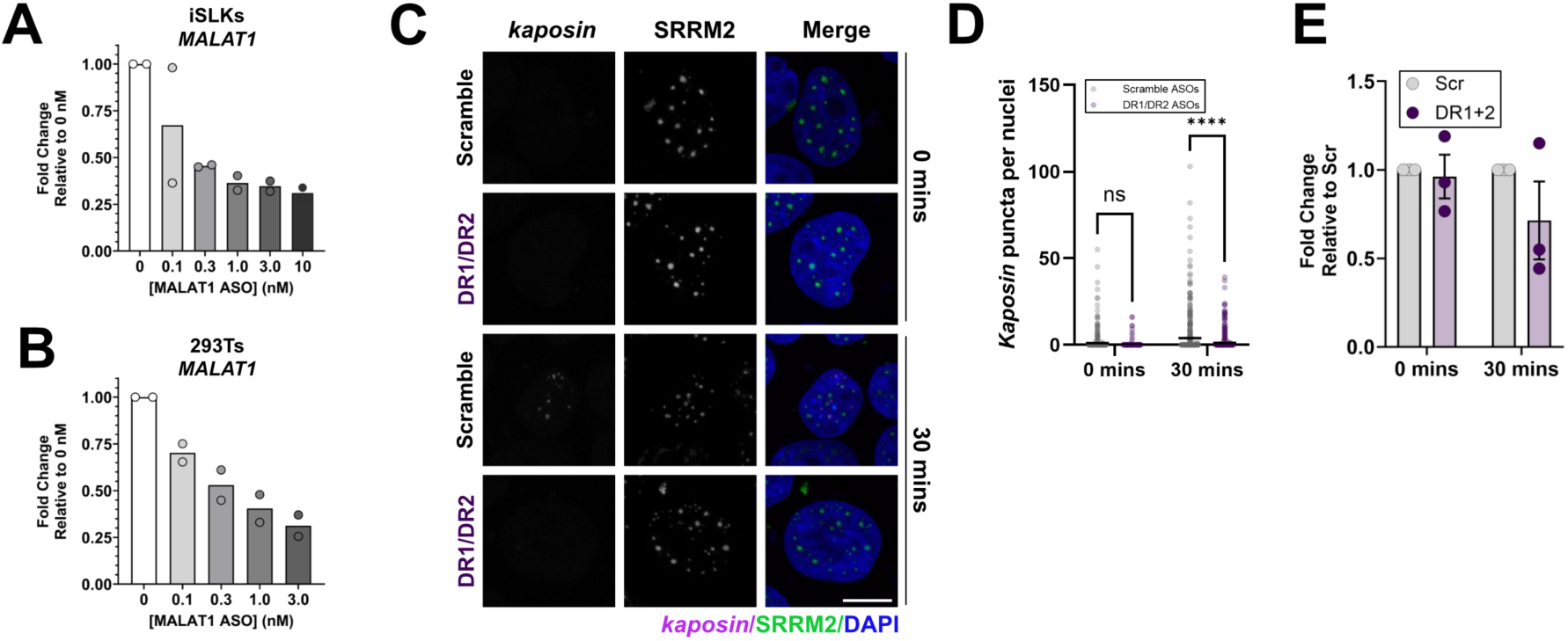
Blocking direct repeat regions prevents *kaposin*-mediated NS seeding. (**A-B**) iSLKs (A) or 293Ts (B) were reverse transfected with the indicated concentrations of a control gapmer ASO and RT-qPCR for *MALAT1* to validate transfection. N=2 (**C-E**) 293T cells were reverse transfected with DR1/DR2 ASOs complementary to *kaposin* or a scrambled control, co-transfected with *kaposinΔstarts and* rtTA constructs and then either stained for kaposin (magenta), SRRM2 (green) and nuclei (blue) (**C-D**) or lysed for RT-qPCR analysis using *kaposin* specific primers (**E**). Statistics were performed using a two-way repeated measures ANOVA test followed by a Tukey’s post-hoc test (N = 3; *p* < 0.05). In D, images were quantified using CellProfiler according to methods.

## Supplemental Video Captions

**Video 1. MCP does not relocalize following doxycycline-inducible *EV-MS2* expression.**

293T cells were transfected with an inducible empty vector construct, an EGFP-tagged MCP (magenta) and rtTA, treated with Dox and then immediately imaged for ∼2 hrs using a Nikon A1R+ confocal microscope. Maximum intensity projections (MIPs) are presented.

**Video 2. MCP relocalizes into large puncta following *kaposinΔstarts-MS2* expression.**

293T cells were transfected with an inducible *kaposinΔstarts* construct, an EGFP-tagged MCP (magenta) and rtTA, treated with Dox and then immediately imaged for ∼2 hrs using a Nikon A1R+ confocal microscope. Maximum intensity projections (MIPs) are presented.

**Video 3. *Kaposin* transcription seeds RBM25-positive NSs.**

293T cells were transfected with an inducible *kaposinΔstarts* construct, an EGFP-tagged MCP (magenta), an mCherry-tagged RBM25 NS marker (green) and rtTA, treated with Dox and then immediately imaged for ∼2 hrs using a Nikon A1R+ confocal microscope. Maximum intensity projections (MIPs) are presented. Images were acquired at 1.5 min interval.

**Video 4. *Kaposin* transcription seeds SRSF1-positive NSs.**

293T cells were transfected with an inducible *kaposinΔstarts* construct, an EGFP-tagged MCP (magenta), an mCherry-tagged SRSF1 NS marker (green) and rtTA, treated with Dox and then immediately imaged for ∼2 hrs using a Nikon A1R+ confocal microscope. Maximum intensity projections (MIPs) are presented. Images were acquired at 3.5 min intervals.

**Video 5. *Kaposin*-induced NSs undergo fusion events.**

293T cells were transfected with an inducible *kaposinΔstarts* construct, an EGFP-tagged MCP (magenta), an mCherry-tagged or SRSF1 (green) and rtTA, treated with Dox and then immediately imaged for ∼2 hrs using a Nikon A1R+ confocal microscope. Maximum intensity projections (MIPs) are presented. Images were acquired at 3.5 min intervals.

